# HIV-1 Hijacks HGF/c-MET Signaling to Promote Viral Entry and Replication in Primary CD4^+^ T Cells

**DOI:** 10.64898/2026.08.03.742552

**Authors:** Shraddha Tripathi, Madeleine M. Allen, Li Wu

## Abstract

The mesenchymal-epithelial transition factor (c-MET) is a receptor tyrosine kinase best known for mediating hepatocyte growth factor (HGF) signaling in cancer, yet its role in HIV-1 infection remains undefined. Here we identify c-MET as a host factor that facilitates HIV-1 replication in CD4^+^ T cells by promoting viral entry. HIV-1 infection upregulates c-MET transcripts and phosphorylation in primary CD4^+^ T cells. Knockout or pharmacological inhibition of c-MET in CD4^+^ T cells significantly reduces HIV-1 Gag expression, viral production, and infectivity. Mechanistically, c-MET disruption reduces cell-surface expression of the HIV-1 receptor CD4 and the coreceptor CXCR4, thereby restricting viral entry and infection. Furthermore, c-MET disruption diminishes activation of NF-κB, STAT1/3, and MAPK signaling pathways that are critical for HIV-1 gene expression. These findings reveal that HIV-1 exploits HGF/c-MET signaling to coordinate entry receptor availability and intracellular signaling, uncovering a previously unrecognized host pathway that supports viral replication in CD4^+^ T cells. Overall, our results highlight an essential role for HGF/c-MET signaling in HIV-1 infection of primary CD4^+^ T cells and suggest a potential host-directed therapeutic target.

**Highlights:**

- c-MET facilitates HIV-1 replication in CD4^+^ T cells by promoting viral entry
- HIV-1 infection upregulates c-MET transcripts and phosphorylation in CD4^+^ T cells
- c-MET disruption reduces CD4 and CXCR4 expression on CD4^+^ T cell surfaces
- c-MET disruption diminishes activation of cellular pathways important for HIV-1

## Introduction

Human immunodeficiency virus type 1 (HIV-1) primarily targets CD4^+^ T cells, leading to progressive immune dysfunction and increased susceptibility to opportunistic infections and malignancies ^1,2^. HIV-1 enters cells through viral envelop protein-mediated binding and fusion to the receptor CD4 and the coreceptor CCR5 or CXCR4 ^3–5^. Despite the success of antiretroviral therapy (ART) in suppressing viral replication, HIV-1 persists as a lifelong infection and global health threat ^1^. The virus manipulates cellular signaling pathways to facilitate replication, evade innate immunity, and induce chronic immune activation ^6,7^. Multiple hosts signaling cascades can be affected by HIV-1, which emerged as an important axis for modulating antiviral responses, promoting immune evasion, and regulating viral replication ^2,6^. Understanding host cell signaling pathways modulated by HIV-1 is essential for developing more effective strategies to combat persistent infection.

Cellular mesenchymal-epithelial transition factor (c-MET), also known as hepatocyte growth factor (HGF) receptor, is a receptor tyrosine kinase expressed in epithelial cells and is frequently dysregulated in multiple cancers ^8–10^. HGF is the ligand for c-MET and primarily secreted by fibroblasts and mesenchymal stromal cells ^11,12^. After tumor-associated fibroblasts produce and secrete inactive precursor HGF, which is converted to active HGF by matriptase and binds to its receptor c-MET ^13^. The binding of the HGF to c-MET results in dimerization of c-MET monomers and auto-phosphorylation of two catalytic tyrosine residues (Y1234 and Y1235) within the kinase activation loop. The subsequent step is phosphorylation of two additional docking tyrosine in the carboxy-terminal tail (Y1349 and Y1356), and when phosphorylated, these tyrosine serves as a docking site for downstream bridging molecules ^14–18^. The binding of downstream docking molecules further activate the downstream pathways of HGF/c-MET, including RAS/MAPK ^19,20^, phosphoinositide 3-kinase / protein kinase B signaling pathway (PI3K/AKT) ^21,22^, Wnt/β-Catenin ^23^, and JAK/STAT ^24^, to drive transcriptome changes and ultimately mediate the phenotypic changes of the cancer cells, including proliferation, migration, invasion, and metastasis ^9,10,25–27^. However, the function of HGF/c-MET has not been reported in the context of HIV-1 infection.

In our previous study, we performed a transcriptomic analysis of 84 type I interferon-responsive genes in peripheral blood mononuclear cells (PBMCs) from viremic people living with HIV-1 (PLWH) and ART-treated individuals, compared with healthy donors ^28^. We found a significant increase in *c-MET* (the gene is also named as *MET* in the literature) mRNA levels in viremic PLWH compared with healthy individuals ^28^. Furthermore, *c-MET* mRNA expression was significantly reduced in PBMCs from HIV-1-suppressed individuals receiving ART compared with viremic PLWH ^28^. Our subsequent transcriptomic analysis of these type I interferon-responsive genes in PBMCs from PLWH with or without cancer also revealed that *c-MET* mRNA expression was significantly elevated in PLWH with cancer compared with PLWH without cancer ^29^. Based on these observations, we sought to characterize the functional significance of c-MET during HIV-1 infection in CD4^+^ T cells.

Here, we report that c-MET is important for HIV-1 infection in primary CD4^+^ T cells by regulating receptor-mediated viral entry. We show that HIV-1 infection upregulates c-MET transcripts and phosphorylation in the HGF-treated primary CD4^+^ T cells and MT-4 cells. Pharmacological inhibition of c-MET in CD4^+^ T cells reduces HIV-1 infection, HIV-1 Gag mRNA and protein levels, viral production, and infectivity. Knockdown (KD) or Knockout (KO) of c-MET in primary CD4^+^ T cells or MT-4 cells results in a decrease in the HIV-1 infection, virion release, and infectivity. Furthermore, c-MET inhibition or KO in CD4^+^ T cells reduces the activation of NF-κB, STAT, and MAPK pathways that are important for efficient HIV-1 gene expression. Mechanistically, c-MET disruption in CD4^+^ T cells decreases the surface expression of CD4 and CXCR4, thereby reducing HIV-1 entry and infection. Overall, our findings demonstrate the pivotal role of HGF/c-MET in regulating HIV-1 replication and host cellular signaling pathways required for viral gene expression in CD4^+^ T cells.

## Results

### HIV-1 infection upregulates c-MET mRNA and phosphorylation of c-MET (p-MET) in primary CD4^+^ T cells and MT-4 cells

We reported that *c-MET* mRNA levels are significantly elevated in PBMCs from viremic PLWH compared with HIV-1-suppressed individuals receiving ART ^28^. We also found that *c-MET* mRNA expression was significantly elevated in PLWH with cancer compared with PLWH without cancer ^29^. An early study reported increased serum HGF concentrations in HIV-1-positive women relative to HIV-1-negative women ^30^. These observations suggest a potential function of HGF/c-MET in regulating HIV-1 infection; however, the role of HGF/c-MET signaling during HIV-1 infection in CD4^+^ T cells remains unexplored. To address this question, we first examined *c-MET* mRNA expression at different time points following HIV-1 infection in primary CD4^+^ T cells isolated from three independent healthy donors. c-MET mRNA levels significantly increased by ∼2.4-fold at 24 h post-infection (hpi), whereas no significant changes were observed at 48 or 72 hpi (Fig. 1A). Similar results were observed in HIV-1-infected MT-4 cells, where *c-MET* mRNA expression increased by ∼2.9-fold at 24 hpi but remained unchanged at later time points (48 and 72 hpi) (Fig. S2A).

**Fig. 1.**
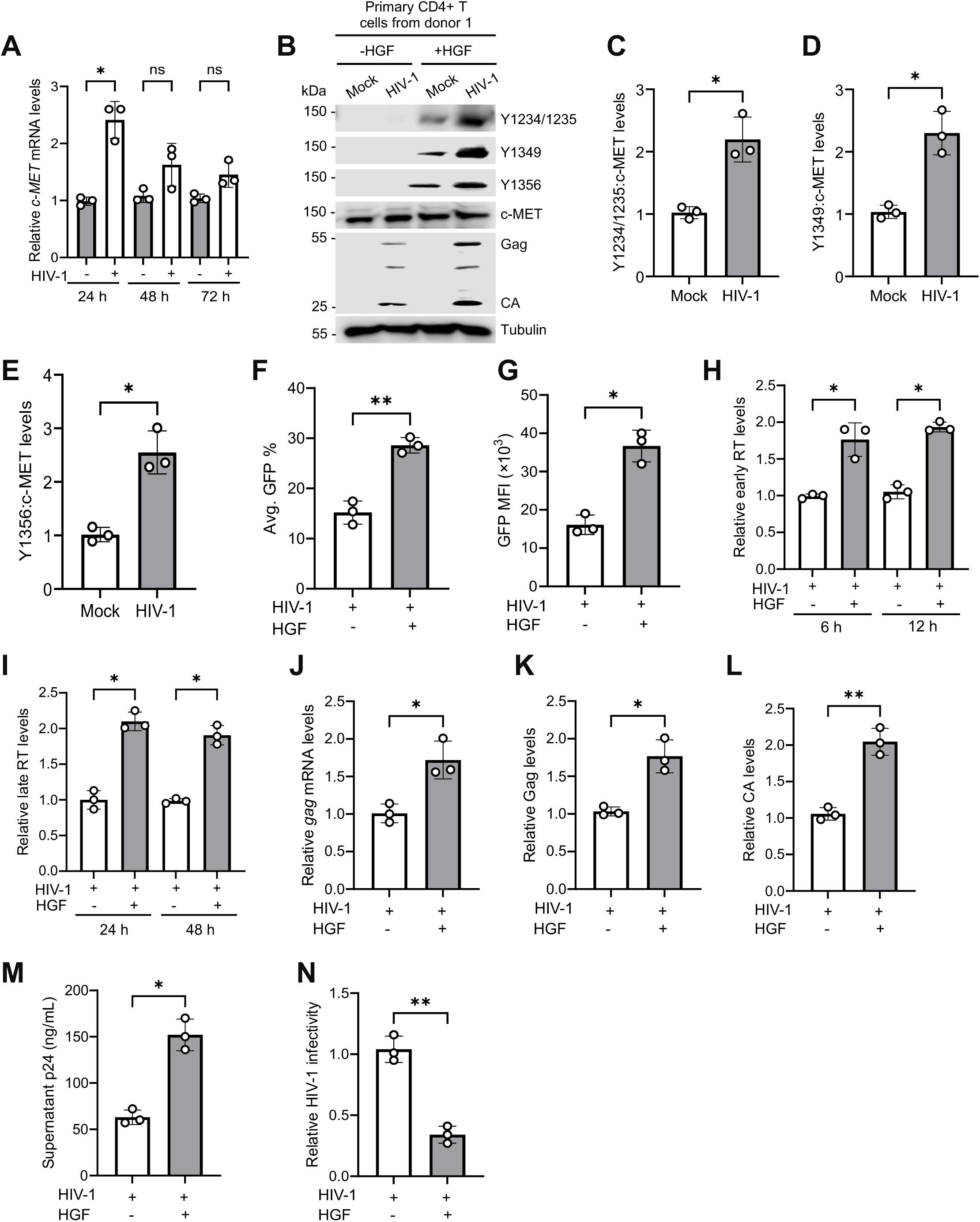
HIV-1 infection promotes HGF-mediated phosphorylation of c-MET in primary CD4^+^ T cells. **(A)** Quantification of c-MET mRNA levels in primary CD4^+^ T cells at 24, 48 and 72 h post HIV-1 infection (hpi) by qRT-PCR. *HPRT* was used as an internal control. Average results from three independent donor’s cells are shown. **(B)** Detection of phosphorylated Y1234/1235, Y1349, and Y1356 of c-MET, total c-MET, HIV-1 Gag and CA, and tubulin by immunoblotting. Tubulin was used as a loading control. Donor 1 results are shown, and the results of donors 2 and 3 are in Fig. S1. **(C-E)** The relative levels of phosphorylated Y1234/1235 (C), Y1349 (D), Y1356 (E) of c-MET were quantified by densitometry analysis and normalized to total c-MET and tubulin. The level of proteins in HGF-treated and mock-infected cells was set to 1. **(F-G)** HIV-1 infection was measured by the expression of GFP reporter using flow cytometry at 48 hpi. Changes in average GFP percentage **(F)** and change in mean fluorescence intensity (MFI) of GFP-positive cells **(G)** from mock or HIV-1 infected primary CD4^+^ T cells from three donors with or without HGF treatment are shown. **(H)** HIV-1 early reverse transcription (early RT) products and **(I)** late RT products were measured by qPCR at the indicated time points in HIV-1-infected cells with or without HGF treatment. Unspliced *GAPDH* was used for normalization. **(J)** HIV-1 *gag* mRNA levels at 48 hpi were quantified using qRT-PCR. *HPRT* was used as an internal control. **(K)** HIV-1 Gag and **(L)** CA protein levels were quantified by densitometry analysis and normalized to tubulin. (K-L) The level of Gag and CA proteins from HIV-1-infected cells without HGF treatment was set to 1. **(M)** HIV-1 p24 levels in the supernatants collected from infected primary CD4^+^ T cells with or without HGF treatment were quantified by ELISA. **(N)** TZM-bl cells were infected with the supernatant (2 ng p24) of HIV-1 infected primary CD4^+^ T cells with or without HGF treatment, and HIV-1 infectivity was measured by luciferase activities at 48 hpi. The level of infected cells without HGF treatment was set to 1. (H-N) Results from three donors’ cells are shown. Paired T-test and one-way ANOVA multiple comparisons test were used to evaluate the statistical significance of the difference between sample groups. * *P* <0.05, ** *P* <0.01, ns, not significant.

Since HGF is the ligand of c-MET and following HGF binding, the kinase activity of MET is switched on by receptor dimerization and phosphorylation at Y1234, Y1235 in the kinase domain, which leads to autophosphorylation of the carboxy-terminal bidentate substrate-binding site (Y1349 and Y1356) and when phosphorylated act as a docking site for the recruitment of many signal-relay molecules ^25,27^. To examine c-MET activation during HIV-1 infection, we treated mock- or HIV-1-infected primary CD4^+^ T cells with or without HGF and then evaluated p-MET levels. We observed no detectable p-MET in the absence of HGF in either mock- or HIV-1-infected primary CD4^+^ T cells (Fig. 1B). We then assessed changes in p-MET in HGF-treated mock- and HIV-1-infected primary CD4^+^ T cells derived from three independent healthy donors (Fig. 1B and S1). HIV-1 infection significantly increased p-MET at tyrosine residues Y1234/1235, Y1349, and Y1356 by ∼2.1-to 2.5-fold in HGF-treated primary CD4^+^ T cells compared with mock-infected and HGF-treated control cells (Fig. 1B–D and S1). A similar increase in p-MET at Y1234/1235, Y1349, and Y1356 was also observed in HGF-treated MT-4 cells following HIV-1 infection relative to mock-infected and HGF-treated cells (Fig. S2B–E). Thus, HIV-1 infection increases the expression of *c-MET* mRNA and HGF-mediated p-MET, suggesting that HIV-1 infection activates the HGF/c-MET signaling in CD4^+^ T cells.

### HGF treatment enhances HIV-1 replication and release but reduces infectivity in CD4^+^ T cells

To assess the effect of HGF treatment on HIV-1 replication, we first measured GFP expression in CD4^+^ T cells infected with a replication-competent, CXCR4-tropic GFP-reporter HIV-1. We observed that HGF treatment significantly increased the average percentage of HIV-1 GFP-positive cells by ∼1.7 fold, along with a ∼2.3-fold increase in the mean fluorescence intensity (MFI) of GFP-positive cells (Fig. 1F–G). Next, we evaluated the effect of HGF on post-entry early events of the HIV-1 lifecycle. We quantified early and late reverse transcription (RT) products at the indicated time points and observed that the levels of early and late RT products significantly increased by ∼1.7-fold and ∼2.0-fold, respectively (Fig. 1H–I). Concordant with the enhanced early and late RT levels, we also observed that *gag* mRNA levels were significantly increased by ∼1.7-fold in HGF-treated cells compared with untreated cells (Fig. 1J). We next assessed HIV-1 Gag protein expression in HIV-1-infected primary CD4^+^ T cells with or without HGF treatment and observed that both HIV-1 Gag, and capsid (CA or p24) protein levels were significantly increased by ∼1.8–2.0-fold following HGF treatment compared with untreated cells (Fig. 1K–L). Furthermore, measurement of p24 release from HIV-1-infected primary CD4^+^ T cells demonstrated that HGF treatment significantly enhanced p24 release by ∼2.4-fold in supernatants derived from three independent healthy donors’ cells compared with cells without HGF treatment (Fig. 1M).

To further evaluate HIV-1 infectivity, we used HeLa-derived TZM-bl cells, which express high levels of CD4, CCR5, and CXCR4 and contain an integrated luciferase reporter gene under the control of the HIV-1 promoter ^31,32^. TZM-bl cells were infected with supernatants containing 2 ng p24-equivalent HIV-1 collected from infected primary CD4^+^ T cells with or without HGF treatment, and luciferase activity was measured at 48 hpi. Unexpectedly, despite the enhanced HIV-1 replication and increased p24 release, HGF treatment significantly reduced viral infectivity in TZM-bl cells by ∼2.4-fold compared with untreated cells (Fig. 1N).

To confirm these observations, we further assessed the levels of p-MET at different tyrosine residues and found a significant increase in p-MET levels at Y1234/1235, Y1349, and Y1356 following HIV-1 infection compared with mock-infected HGF-treated MT-4 cells (Fig. S2B–E). HGF treatment significantly enhanced HIV-1 replication, as evidenced by increased the percentage of HIV-1-GFP-positive cells and GFP expression levels (Fig. S2F-G), elevated levels of early and late RT products (Fig. S2H-I), increased *gag* mRNA expression (Fig. S2J), higher Gag and CA protein expression (Fig. S2K-L), and enhanced p24 release (Fig. S2M). We next evaluated HIV-1 infectivity in TZM-bl cells by infecting the cells with 2 ng p24-equivalent virus collected from the supernatants of infected MT-4 cells with or without HGF treatment. Consistently, despite the enhanced HIV-1 replication, HGF treatment significantly reduced HIV-1 infectivity of virions released from HGF treated primary CD4^+^ T cells or MT-4 cells in TZM-bl cells (Fig. S2N). Together, these results indicate that HGF treatment enhances HIV-1 replication and p24 release in CD4^+^ T cells but unexpectedly reduces viral infectivity in TZM-bl cells.

### Productive HIV-1 infection is required for p-MET upregulation in CD4^+^ T cells

To determine whether the increase in p-MET is a direct consequence of HIV-1 infection, we used pharmacological inhibitors targeting two distinct stages of the viral replication cycle. AMD3100 (AMD) was used to block viral entry/fusion, while nevirapine (NVP) was used to inhibit reverse transcription. Primary CD4^+^ T cells from three independent healthy donors were pretreated with HGF and AMD or NVP for 1 h, followed by mock or HIV-1 infection for 48 h, and DMSO was used as a vehicle control. AMD treatment significantly reduced HIV-1–induced p-MET at Y1234/1235 (Fig. 2A–B, S3A–B), Y1349 (Fig. 2A, 2C, S3A–B), and Y1356 (Fig. 2A, 2D, S3A–B) by approximately 1.5-to 3.2-fold, respectively, compared with control cells. Similarly, NVP treatment also markedly decreased HIV-1–induced p-MET at Y1234/1235 (Fig. 2E–F, S3A–B), Y1349 (Fig. 2E, 2G, S3A–B), and Y1356 (Fig. 2E, 2H, S3A–B) by approximately 2-to 3-fold compared with control cells. We further assessed p-MET levels in MT-4 cells following the antiviral treatment. Consistently, we observed a similar decrease in the HIV-1 induced p-MET at Y1234/1235, Y1349 and Y1356 residues in MT-4 cells treated with either AMD or NVP (Fig. S4). These results show that inhibiting HIV-1 replication at either the entry or reverse transcription stage reduces p-MET expression, indicating that productive HIV-1 infection is required for the increase in p-MET levels in CD4^+^ T cells.

**Fig. 2.**
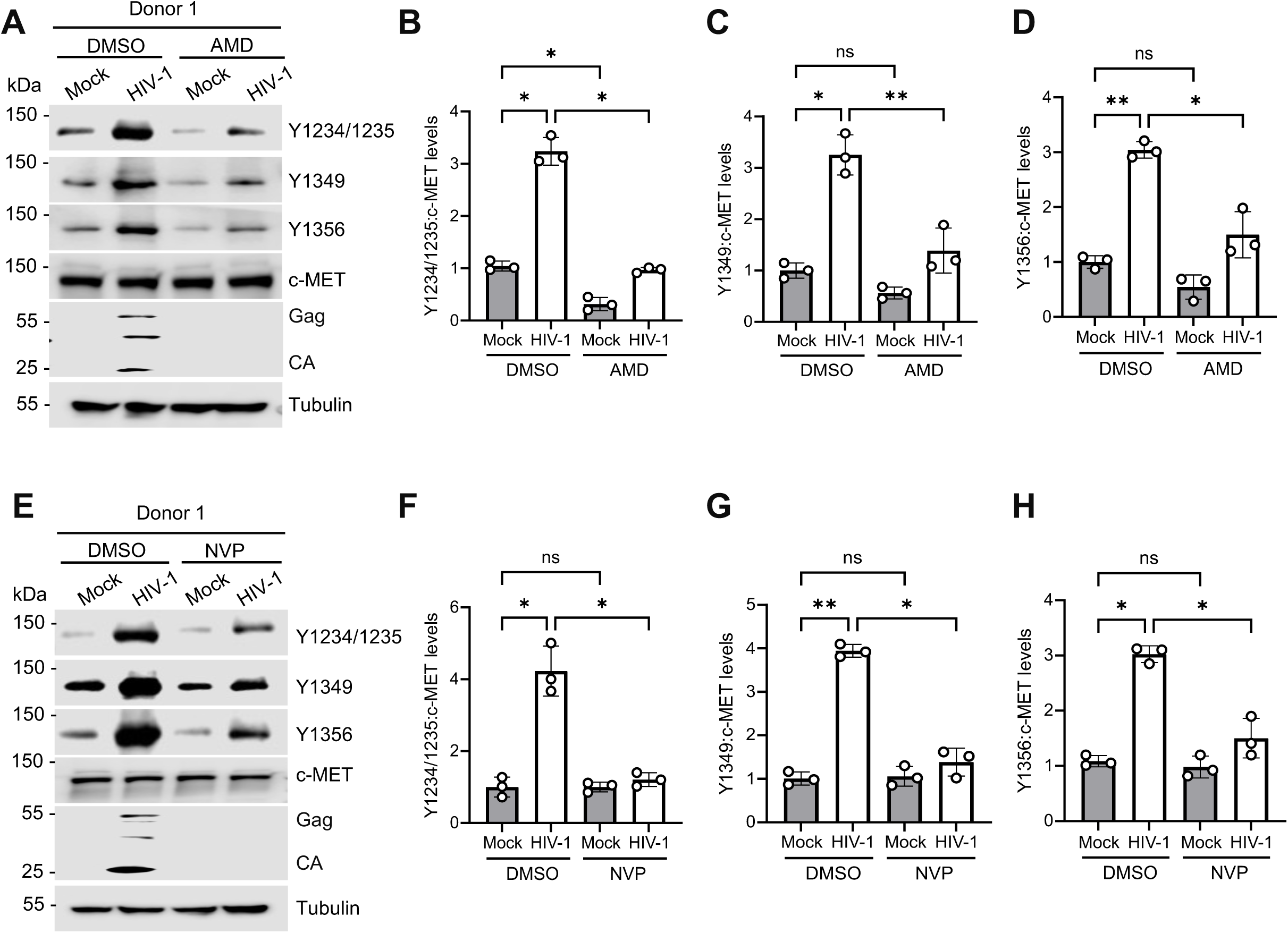
AMD3100 (AMD) or nevirapine (NVP) treatment decreases phosphorylation of c-MET in primary CD4^+^ T cells during HIV-1 infection. **(A and E)**. Detection of phosphorylated Y1234/1235, Y1349, and Y1356 of c-MET, total c-MET, HIV-1 Gag, CA, and tubulin by immunoblotting in mock or HIV-1-infected cells treated with DMSO (control) or AMD **(A)** or NVP **(E)**. Tubulin was used as a loading control. Donor 1 results are shown, and results of donors 2 and 3 are in Fig. S3. The relative levels of Y1234/1235 **(B and F)**, Y1349 **(C and G)**, Y1356 **(D and H)** in cells treated with AMD **(B-D)** or NVP (**F-H)** were quantified by densitometry analysis and normalized to total c-MET and tubulin. The level of proteins in DMSO-treated and mock-infected cells was set to 1. Two-way ANOVA, Šídák’s multiple comparisons test was used to evaluate the statistical significance of the difference between sample groups. * *P* <0.05, ** *P* <0.01, ns, not significant.

### Treatment with a c-MET inhibitor decreases HIV-1 replication in CD4^+^ T cells

To investigate the role of HGF/c-MET signaling in HIV-1 replication, we inhibited c-MET activation in MT-4 cells during HIV-1 infection using the c-MET inhibitor, Crizotinib (PF-2341006), which was developed for cancer treatment in clinic ^33–35^. Primary CD4^+^ T cells are isolated from three independent healthy donors were pretreated with HGF, crizotinib, or DMSO vehicle control for 1 h prior to HIV-1 infection. We first measured p-MET levels at three tyrosine residues and found that crizotinib treatment significantly reduced phosphorylation at Y1234/1235, Y1349, and Y1356 (Fig. 3A-D and S5).

**Fig. 3.**
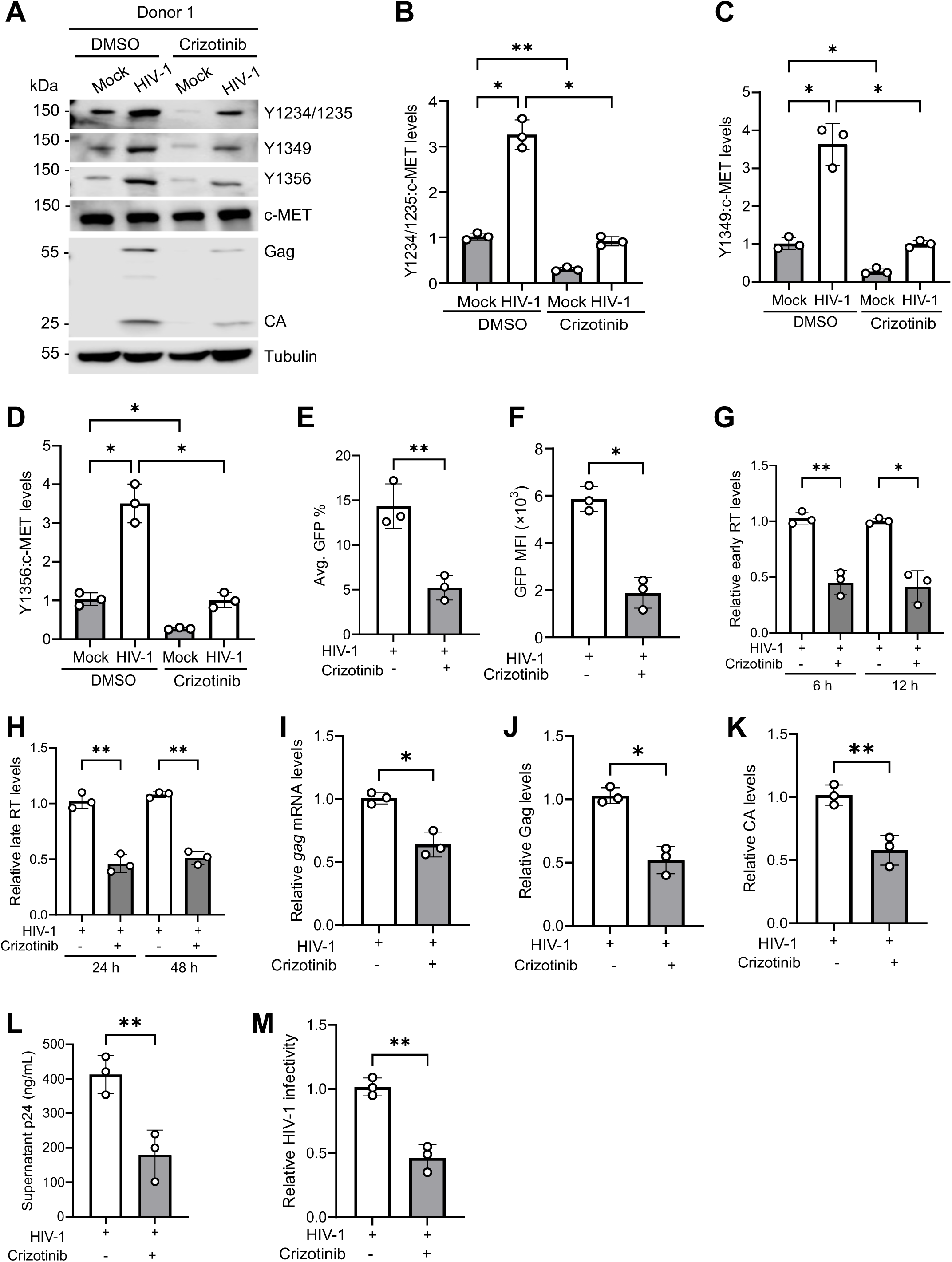
Crizotinib treatment decreases HIV-1 replication and infectivity in primary CD4^+^ T cells. **(A)** Detection of phosphorylated Y1234/1235, Y1349, and Y1356 of c-MET, total c-MET, HIV-1 Gag, CA, and tubulin by immunoblotting in mock or HIV-1 infected primary CD4^+^ T cells treated with DMSO or crizotinib. Tubulin was used as a loading control. Results of donor 1 are shown and the results of donors 2 and 3 are in Fig. S5. **(B-D)** The relative levels of Y1234/1235 (B), Y1349 (C), Y1356 (D) were quantified by densitometry analysis and normalized to total c-MET and tubulin. The level of proteins in mock-infected HGF treated cells was set to 1. **(E-F)** HIV-1 infection was measured by GFP expression at 48 hpi. Changes in average GFP percentage (E) and MFI of GFP-positive cells (F) from HIV-1 infected cells treated with DMSO or crizotinib. **(G)** HIV-1 early RT products and **(H)** late RT products were measured by qPCR at the indicated time points in HIV-1 infected primary CD4^+^ T cells treated with DMSO or without crizotinib. Unspliced *GAPDH* was used for normalization. **(I)** HIV-1 *gag* mRNA levels in cells treated with DMSO or crizotinib were quantified using qRT-PCR. The amplification of *HPRT* was used as an internal control. **(J-K)** HIV-1 Gag and CA levels were quantified by densitometry analysis and normalized to tubulin. The level of Gag (J) and CA (K) proteins from HIV-1 infected DMSO treated cells was set to 1. **(L)** HIV-1 p24 levels in the supernatants collected from HIV-1 infected primary CD4^+^ T cells treated with DMSO or Crizotinib were quantified by ELISA. **(M)** TZM-bl cells were infected with the supernatant of HIV-1-infected primary CD4^+^ T cells treated with DMSO or crizotinib, and HIV-1 infectivity was measured by luciferase activities at 48 hpi. The level of infected cells with DMSO treatment was set to 1. (F-M) Results from three donors’ cells are shown. Paired T-test and one-way ANOVA multiple comparisons test were used to evaluate the statistical significance of the difference between sample groups. * *P* <0.05, ** *P* <0.01.

To assess the role of p-MET in HIV-1 replication in primary CD4⁺ T cells, we first examined the effect of crizotinib on HIV-1 GFP expression. Crizotinib treatment significantly decreased the percentage of GFP-positive cells by ∼2.7-fold (Fig. 3E) and the GFP levels by ∼3-fold (Fig. 3F). Next, we assessed the levels of HIV-1 early and late RT products following crizotinib treatment in primary CD4^+^ T cells. We observed that both early and late RT products were significantly reduced upon crizotinib treatment at 6-48 hpi (Fig. 3G-H). Compared with control cells, we observed that HIV-1 *gag* mRNA levels were significantly decreased by ∼1.5-fold following crizotinib treatment of primary CD4^+^ T cells from three independent healthy donors (Fig. 3I). We further evaluated HIV-1 Gag and CA/p24 protein expression. Notably, crizotinib treatment resulted in ∼1.9-fold and ∼2.3-fold decreases in HIV-1 Gag and CA protein levels, respectively, compared with DMSO-treated and HIV-1-infected cells at 48 hpi (Fig. 3J–K). We next examined p24 release from HIV-1-infected primary CD4^+^ T cells treated with either crizotinib or DMSO. Crizotinib treatment significantly reduced p24 release by ∼2.2-fold compared with DMSO-treated cells (Fig. 3L). To further assess viral infectivity, HIV-1 input was normalized based on p24 content prior to infection of TZM-bl cells. The results showed that the infectivity of HIV-1 produced from crizotinib-treated CD4^+^ T cells was significantly reduced by ∼2.2-fold compared with virus produced from DMSO-treated cells (Fig. 3M). These findings suggest that HGF/c-MET signaling positively regulates HIV-1 replication in CD4^+^ T cells.

Consistent with the findings in primary CD4^+^ T cells, we observed a significant reduction in p-MET at Y1234/1235, Y1349, and Y1356 in both mock- and HIV-1-infected MT-4 cells following crizotinib treatment compared with DMSO-treated cells (Fig. S6A–D). Crizotinib treatment also significantly reduced the percentage of HIV-1 GFP-positive cells and the GFP levels (∼2.6-fold), HIV-1 early and late RT products (∼2-fold), *gag* mRNA levels (∼1.6-fold), and Gag and CA protein expression (∼1.5-fold) in MT-4 cells compared with HIV-1-infected DMSO-treated cells (Fig. S6E–K). Concordantly, p24 release and HIV-1 infectivity were also significantly decreased by ∼2-fold in crizotinib-treated MT-4 cells compared with control cells (Fig. S6L–M). Thus, inhibition of c-MET activity leads to efficient downregulation of HIV-1 infection, viral protein expression, release and infectivity.

### Crizotinib treatment reduces activation of the NF-κB, STAT, and MAPK signaling in CD4^+^ T cells

HIV-1 infection is known to regulate the NF-κB, JAK/STAT and p38 MAPK signaling pathways in CD4^+^ T cells ^36–41^. Previous studies have shown that HGF/c-MET pathway initiates multiple downstream signaling cascades, including the Erk/MAPK, JAK-STAT, and NF-κB signaling pathways ^9,10,20,42,43^. NF-κB activation is a hallmark of HIV-1 infection and is stimulated by multiple viral components, including HIV-1 RNA and proteins in CD4^+^ T cells ^44–48^. To investigate the potential role of c-MET in regulating NF-κB signaling, we examined the effect of crizotinib on the NF-κB signaling proteins. We measured the effect of crizotinib treatment on the key components of the NF-κB signaling pathway in primary CD4^+^ T cells from three independent healthy donors. Compared with the mock-infected and HGF-treated cells, the NF-кB activation markers, phosphorylated IKKα/β (p-IKKα/β) and phosphorylated IKBα (p-IKBα) protein levels, were significantly increased in the HIV-1 infected and HGF-treated cells, and crizotinib treatment significantly attenuated HIV-1 induced p-IKKα/β (Fig. 4A–B and S7A) and p-IKBα (Fig. 4A, C and S7A) to the basal levels compared with the DMSO and HGF-treated cells.

**Fig. 4.**
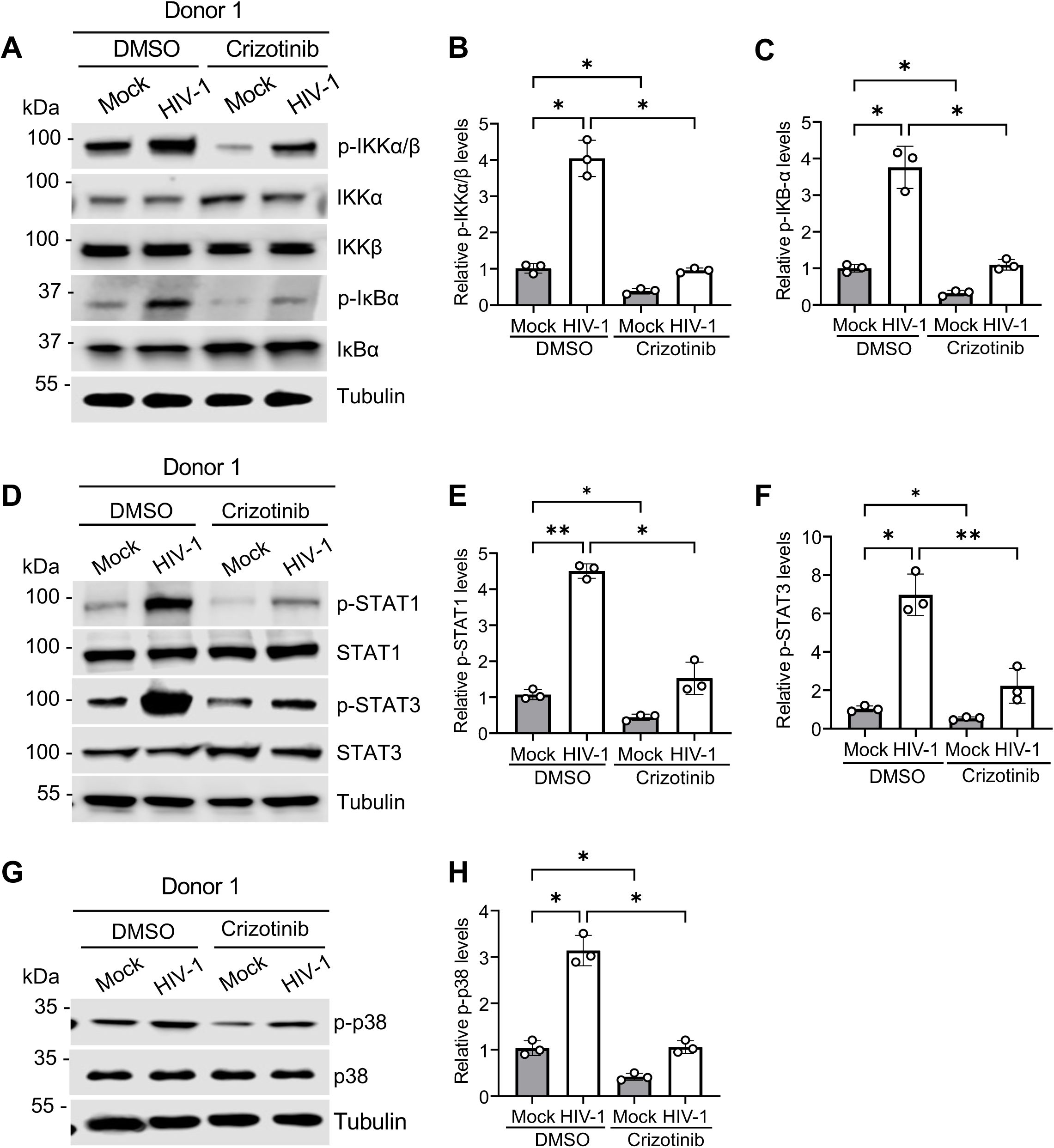
Crizotinib treatment reduces p-IKKα/β, IκBα, p-STAT1, p-STAT3, and p-p38 MAPK in primary CD4^+^ T cells during HIV-1 infection. **(A, D, and G)** Mock or HIV-1 infected primary CD4^+^ T cells were treated with DMSO or crizotinib. The cell lysates were harvested and (A) p-IKKα/β, IKKα, IKKβ, p-IκBα, IκBα, and tubulin (D) p-STAT1, STAT1, p-STAT3, STAT3, tubulin, and (G) p-p38, p38, and tubulin were detected by immunoblotting. Tubulin was a loading control. Results from donor 1 are shown and results of donors 2 and 3 are in Fig. S7. **(B, C, E, F, H)** The relative p-IKKα/β (B), p-IκBα (C), p-STAT1, (E) p-STAT3 (F), and p-p38 (H) levels were quantified by densitometry analysis. Relative levels were normalized to tubulin. The level of proteins in mock-infected DMSO treated cells was set to 1. Two-way ANOVA, Šídák’s multiple comparisons test was used to evaluate the statistical significance of the difference between sample groups. * *P* <0.05, ** *P* <0.01.

Activation of c-MET leads to phosphorylation of STAT proteins (p-STAT), promoting their dimerization and nuclear translocation, where they regulate the transcription of genes involved in cell survival, proliferation, and inflammation ^24,49–52^. To better understand the effect of c-MET activation on STAT signaling, we investigated the effect of the crizotinib treatment on JAK/STAT signaling proteins, particularly STAT1 and STAT3. To this end, we analyzed p-STAT1, total STAT1, p-STAT3, and total STAT3 levels following crizotinib treatment and HIV-1 infection. Compared with mock-infected MT4 cells, HIV-1 infection significantly increased p-STAT1 and p-STAT3 levels (Fig. 4D–F and S7B). Furthermore, inhibition of c-MET by crizotinib markedly reduced p-STAT1 (∼3-fold) and p-STAT3 (∼4-fold) expression respectively, compared with DMSO controls (Fig. 4D–F and S7B).

HGF/c-MET signaling is known to regulate the p38 MAPK signaling pathway in different cancers ^20,53^. Previous studies have shown that HIV-1 infection of both primary CD4^+^ T cells and different T cell lines rapidly activated the cellular p38 MAPK pathway to enhance viral replication ^36,37^. Since both HIV-1 and HGF/c-MET are known to modulate p38 MAPK signaling, we investigated the effect of the crizotinib treatment on phosphorylated-p38 MAPK (p-p38) and total p38 proteins. Compared with mock-infected cells, HIV-1 infection significantly increased p-p38 MAPK levels, while inhibition of c-MET by crizotinib markedly reduced p-p38 expression by ∼3-fold in primary CD4^+^ T cells from three independent healthy donors (Fig. 4G–H and S7C). Consistently, crizotinib treatment also resulted in the significant downregulation of the p-IKBα, p-IKKα/β, p-STAT1, p-STAT3 and p-p38 expression in mock or HIV-1-infected MT-4 cells compared with the DMSO-treated MT-4 cells (Fig. S8). These results suggest that c-MET modulates HIV-1 replication and gene expression through the regulation of NF-кB, JAK/STAT and p38 MAPK signaling in CD4^+^ T cells.

### c-MET knockdown (KD) in primary CD4^+^ T cells reduces HIV-1 replication and infectivity

To evaluate the role of c-MET in regulating HIV-1 replication in primary CD4^+^ T cells, we used nucleofection to deliver two independent CRISPR RNAs (crRNAs) targeting *c-MET* (indicated as KD-1 and KD-2). KD of endogenous c-MET was confirmed by immunoblotting (Fig. 5A and S9). We first examined the HIV-1-GFP infection by measuring the GFP expression using flow cytometry. The results showed that both c-MET KD-1 and KD-2 significantly reduced both the percentage and MFI of GFP-positive cells compared with control cells (Fig. 5B–C). We then assessed *gag* mRNA and protein expression in control and c-MET KD cells, *gag* mRNA expression levels were significantly downregulated by ∼2-fold in the c-MET KD cells as compared with the control cells (Fig. 5D). c-MET KD also significantly decreased Gag expression (∼2-fold) and CA protein (∼2.5-fold) expression in both KD-1 and KD-2 cells compared with control cells (Fig. 5E–F and S9). Furthermore, HIV-1 p24 release in cell supernatants was significantly decreased in the c-MET KD cells compared with the control cells (Fig. 5G). Moreover, HIV-1 input was normalized by 2 ng of p24 content prior to infection of TZM-bl indicator cells to measure viral infectivity. The results confirmed that the infectivity of HIV-1 produced from c-MET KD cells was significantly decreased by 2-fold compared with the viruses from the control cells (Fig. 5H). These results suggest an important role of endogenous c-MET in facilitating HIV-1 replication in primary CD4^+^ T cells.

**Fig. 5.**
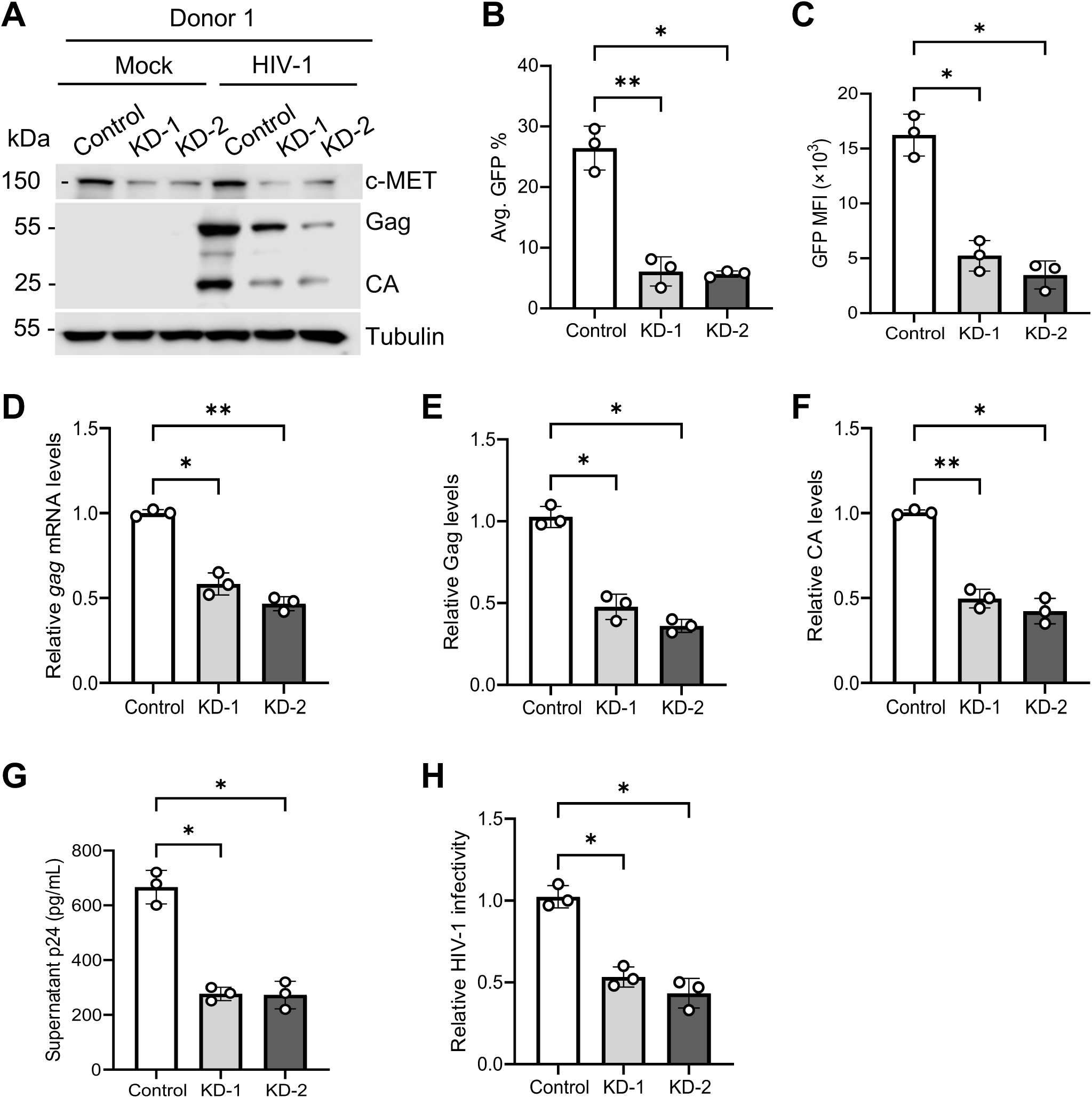
KD of c-MET in primary CD4^+^ T cells reduces HIV-1 replication and infectivity. **(A-H)** KD of c-MET expression in activated primary CD4^+^ T cells from three donors were with CRISPR-Cas9 ribonucleoprotein (crRNP) targeting two different sites in *c-MET* (KD-1 and KD-2). **(A)** Expression of c-MET and HIV-1 Gag, CA, and tubulin was detected by immunoblotting. Tubulin was a loading control. Results of donor 1 are shown. Results of donor 2-3 are in Fig. S9. **(B-H)** Three dots on each bard represent the results of three donors. **(B-C)** HIV-1 infection was measured by GFP expression using flow cytometry at 48 hpi in HIV-1-infected control and c-MET KD cells. Changes in average GFP percentage (B) and MFI of GFP-positive cells (C) from HIV-1 infected control or c-MET KD cells of three donors. **(D)** HIV-1 *gag* mRNA levels in control and c-MET KD cells were quantified using qRT-PCR and *HPRT* was used as an internal control. **(E-F)** The relative Gag (E) and CA (F) levels were quantified by densitometry analysis. Relative Gag and CA levels were normalized to tubulin. The level of proteins in HIV-1-infected control cells was set to 1. **(G)** HIV-1 p24 levels in the supernatants from infected control and c-MET KD cells were quantified by ELISA. **(H)** TZM-bl cells were infected with the supernatants (2 ng of p24) from HIV-1 infected control and c-MET KD cells, and HIV-1 infectivity was measured by luciferase activities at 48 hpi. The level of HIV-1-infected control cells was set to 1. One-way ANOVA multiple comparisons test was used to evaluate the statistical significance of the difference between sample groups. * *P* <0.05, ** *P* <0.01.

### c-MET KO in MT-4 cells reduces HIV-1 replication and infectivity

To understand the mechanism by which c-MET regulates HIV-1 infection, we generated c-MET KO MT-4 cells based on the lentiCRISPRv2 system and two different *c-MET*-specific single guide RNAs (sgRNA). The empty vector that expresses negative control sgRNA was used to generate control cells. MT-4 cells transduced with the lentiviral vectors were selected by puromycin and clonal purified with limiting dilution. We obtained single clones of the respective cell populations treated with sgRNA1 (KO-1) or sgRNA2 (KO-2) having negligible expression of c-MET compared with the control cells (Fig. 6A). These stable cell clones were used for the subsequent study.

**Fig. 6.**
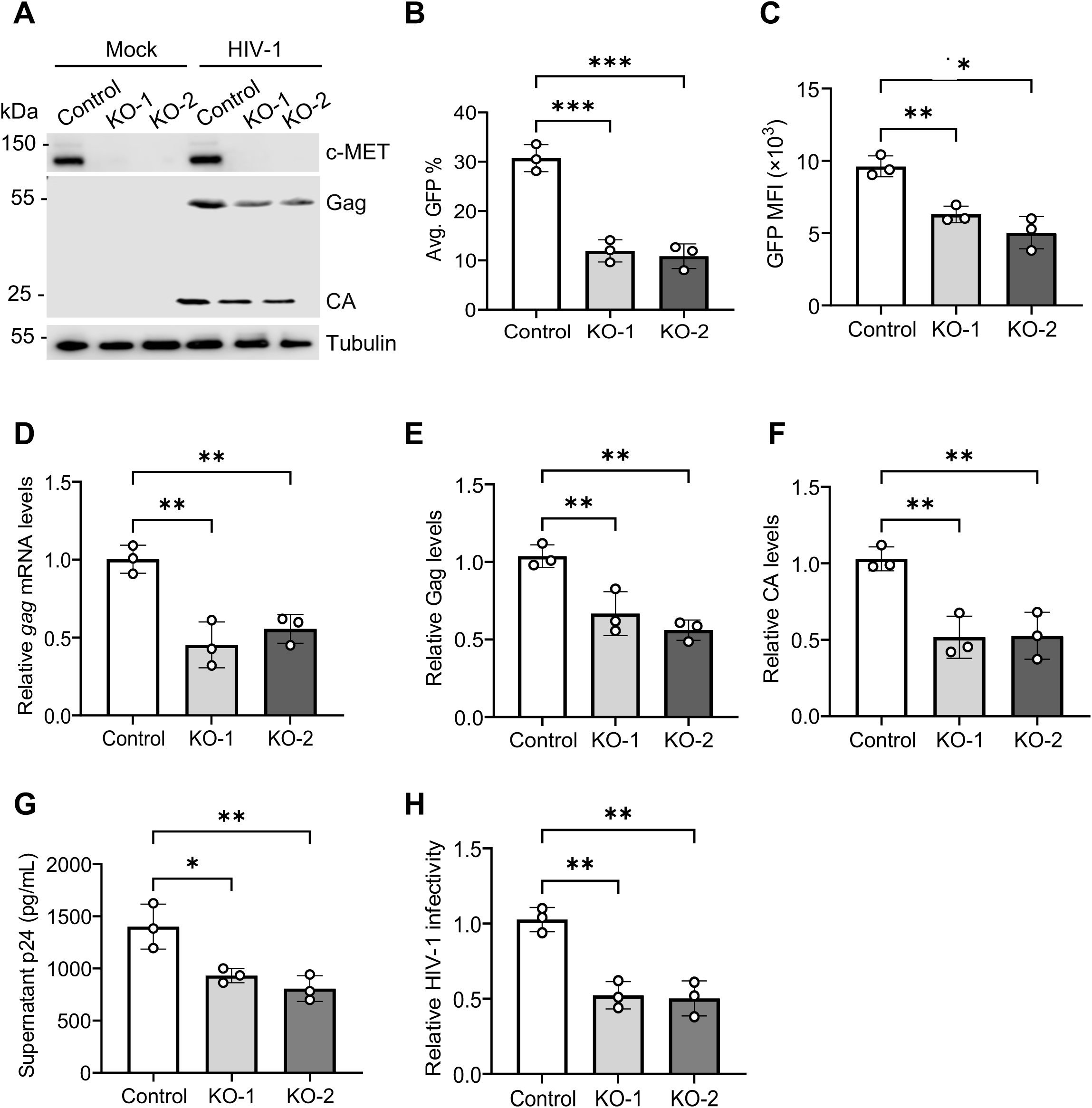
KO of c-MET in MT-4 cells decreases HIV-1 replication and infectivity. **(A)** Control and c-MET KO MT-4 cells were mock or HIV-1 infected. The cell clone was obtained from guide RNA 1 (KO-1) and guide RNA 2 (KO-2) treated population MT-4 cells. c-MET, Gag, CA, and tubulin in cell lysates were detected by immunoblotting. Tubulin was a loading control. (**B-C)** HIV-1 infection was measured by GFP expression using flow cytometry at 48 hpi in HIV-1 infected control and c-MET KO MT-4 cells. Changes in average GFP percentage (B) and MFI of GFP-positive cells (C) from HIV-1 infected control or c-MET KO MT-4 cells. **(D)** HIV-1 *gag* mRNA levels in control and c-MET KO-1 MT-4 cells were quantified using qRT-PCR and *HPRT* was used as an internal control. **(E-F)** The relative Gag (E) and CA (F) levels were quantified by densitometry analysis. Relative Gag and CA levels were normalized to tubulin. The level of proteins in HIV-1-infected control MT-4 cells was set to 1. **(G)** HIV-1 p24 levels in the supernatants from infected control and c-MET KO cells were quantified by ELISA. **(H)** TZM-bl cells were infected with the supernatants (2 ng of p24) from HIV-1 infected control and c-MET KO MT-4 cells, and HIV-1 infectivity was measured by luciferase activities at 48 hpi. The level of HIV-1-infected control MT-4 cells was set to 1. One-way ANOVA multiple comparisons test was used to evaluate the statistical significance of the difference between sample groups. * *P* <0.05, ** *P* <0.01.

We first examined the HIV-1-GFP infection by measuring the GFP expression using flow cytometry. The results showed that c-MET KO significantly reduced both the percentage (∼2.1-to ∼2.8-fold) and MFI of GFP-positive cells (∼2-to ∼2.5-fold) compared with control cells (Fig. 6B–C). Next, we assessed *gag* mRNA and protein expression in control and c-MET KO cells, *gag* mRNA expression levels were significantly downregulated by ∼2-to 2.2-fold in the c-MET KO-1 KO-2 cells as compared with the control cells (Fig. 6D). c-MET KO also decreased Gag expression (∼1.5-to ∼2-fold) and CA protein expression (∼2-to ∼2.2-fold) in both KO-1 and KO-2 cells compared with control cells (Fig. 6A, 6E–F). Furthermore, we measured HIV-1 p24 release in cell supernatants and found significant decrease in the c-MET KO cells compared with the control cells (Fig. 6G). Moreover, HIV-1 input was normalized by 2 ng of p24 content prior to infection of TZM-bl indicator cells to measure viral infectivity. The results confirmed that the infectivity of HIV-1 produced from c-MET KO cells was significantly decreased by 2-fold compared with the viruses from the control cells (Fig. 6H). Thus, c-MET KO in MT-4 cells significantly reduces HIV-1 infection, viral production, and infectivity, suggesting an important role of c-MET in facilitating HIV-1 replication in CD4^+^ T cells.

### c-MET KO in MT-4 cells reduces activation of the NF-κB, STAT, and MAPK signaling

To examine the effect of c-MET KO on the signaling pathways that are important for HIV-1 gene expression, we measured the NF-κB, STAT1/3 and p-38 MAPK signaling proteins during HIV-1 infection. We found that c-MET KO decreased HIV-1-induced p-IKKα/β levels by ∼4-fold in KO-1 and ∼4.2-fold in KO-2 cells compared with control cells (Fig. S10A–B). We also observed that p-IKBα levels were significantly downregulated by ∼4.6-to 4.8-fold in c-MET KO-1 and KO-2 cells compared with the control cells (Fig. S10A and C). We next measured p-STAT1 levels in c-MET KO MT-4 cells and we observed a significant decrease by ∼3.5-to 3.6-fold compared with the control cells (Fig. S10D–E). Similarly, p-STAT3 levels were significantly downregulated by ∼5.4-fold in c-MET KO cells compared with the control cells (Fig. S10D and F). Furthermore, the levels of p-p38 MAPK were also significantly reduced by ∼2.5-to 2.8-fold in c-MET KO cells compared with the control cells (Fig. S10G–H). These results confirm that endogenous c-MET in CD4^+^ T cells is important for activation of the NF-κB, STAT1/3, and p-38 MAPK signaling pathways upon HIV-1 infection.

### c-MET KO or crizotinib treatment of CD4^+^ T cells reduces CD4 and CXCR4 expression

To gain mechanistic insights into how c-MET KO suppressed HIV-1 replication, we performed RNA sequencing to understand the whole transcriptomic profiles of MT-4 control, KO-1, and KO-2 cells using biological triplicates of RNA samples from each cell type. Next, by analyzing the differential expressed genes (DEGs) implicated in both KO-1 and KO-2 having more than 2-fold change (FC) and with false discovery rate (FDR)-adjusted *P* values□<□0.05, we found that 6,446 genes were significant DEGs in control vs. c-MET KO cells. Among these DEGs, 3,458 were significantly upregulated and 2,988 were significantly downregulated in c-MET KO cells as compared with control MT-4 cells (Supplemental Excel file). Top 30 DEGs are shown as a representative in the heat map (Fig. 7A). To identify the pathways related to cellular transcripts that are regulated upon c-MET KO, we conducted gene ontology (GO) pathway analysis (Fig. 7B). Top 20 GO terms with the lowest *P* values were plotted on a bubble plot against the gene ratio (i.e. number of DEGs per all annotated genes of the GO term). The analysis indicates that c-MET significantly regulates several pathways, including TNF-α signaling through NF-κB, interferon-γ response, epithelial mesenchymal response, unfolded protein response, and inflammatory response (Fig. 7B).

**Fig. 7.**
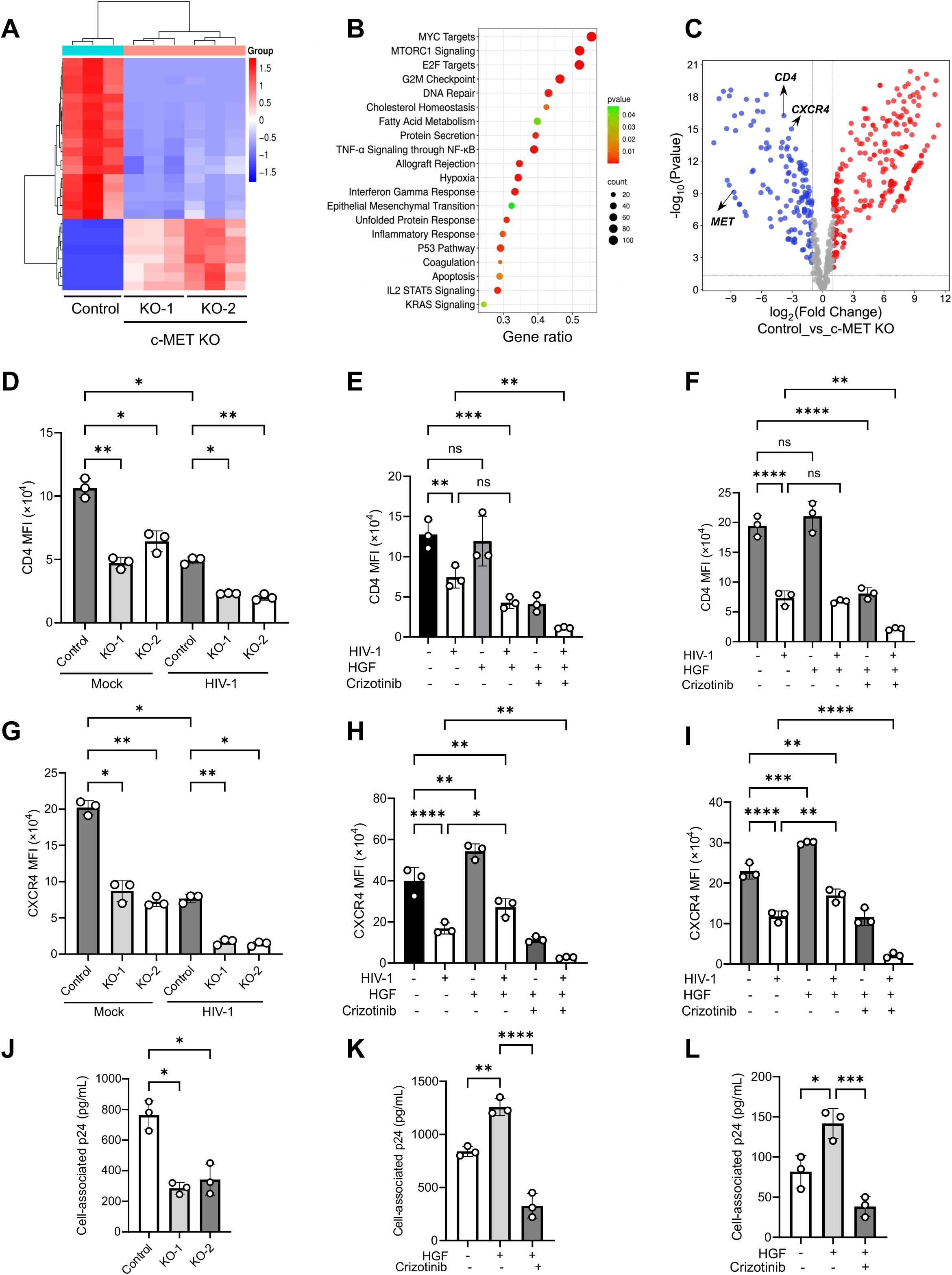
c-MET KO in MT-4 cells or crizotinib treatment in CD4^+^ T cells decreases HIV-1 entry and infection by downregulating CD4 and CXCR4 expression levels. **(A)** Heat map showing transcript-level differences between control and c-MET KO MT-4 cells. Due to the large data set, only 30 genes are displayed. Each row represents a transcript, and each column represents a sample. Both rows and columns are clustered using correlation distance. Up-regulated and down-regulated genes are shown in red and blue, respectively. The numbers next to the figure represent raw Z-scores. **(B)** The differentially expressed genes (DEGs) are obtained from control and c-MET KO MT-4 cells were used for gene ontology enrichment analysis. Top 20 GO terms with the lowest p values were plotted on a bubble plot against the gene ratio (i.e. number of DEGs per all annotated genes of the GO term). Significance is indicated by sphere color. The size of the sphere corresponds to the number of DEGs associated with each pathway. **(C)** Volcano plot showing DEGs from control vs c-MET KO MT-4 cells with p value against log2 fold change. The known HIV-1 entry receptor *CD4* and co-receptor *CXCR4* as well as *MET* were identified as downregulated genes. The relative MFI of CD4 **(D)** and CXCR4 **(G)** positive cells were determined by flow cytometry in mock or HIV-1 infected control and c-MET KO MT-4 cells. The relative MFI of CD4 **(E-F)** and CXCR4 **(G-H)** positive cells were determined by flow cytometry upon treatment with either HGF, DMSO or crizotinib in mock or HIV-1-infected primary CD4^+^ T cells from 3 different donors (E and H) and MT-4 cells (F and I). **(J-L)** Cell-associated HIV-1 p24 was measured in control and c-MET KO MT-4 cells (J), primary CD4^+^ T cells (K), and MT-4 cells (L) after incubation with HIV-1 for 2 h at 37°C. After extensive washes, cells were trypsinized and lysed for p24 measurement by ELISA. Two-way ANOVA, Šídák’s multiple comparisons test was used to evaluate the statistical significance of the difference between sample groups. * *P* <0.05, ** *P* <0.01, *** *P* <0.001, **** *P* <0.0001, ns, not significant.

A volcano plot illustrates the distribution of the significantly differentially expressed mRNAs ranked by log₂ FC (Fig. 7C). Volcano plot showing upregulated (red) and downregulated (blue) mRNA from c-MET KO MT-4 cells compared with the control (FC ≥ 2-fold, *P* < 0.05). Interestingly, among the differentially expressed genes, the HIV-1 receptor and coreceptor, *CD4* and *CXCR4* genes were significantly downregulated upon the c-MET KO cells (Fig. 7C, the c-*MET* gene is shown as a control). To validate the downregulation of the CD4 and CXCR4, we first measured the transcript level using quantitative RT-PCR (qRT-PCR) and found significant downregulation of CD4 by ∼2 FC in KO-1 and ∼2.9 FC in KO-2 MT-4 cells as compared with control cells (Fig. S11A). We also measured the levels of CD4 transcript levels upon crizotinib inhibitor treatment, and we observed that it significantly reduces the *CD4* transcript levels in primary CD4^+^ T cells by ∼2 FC (Fig. S11B) and in MT-4 cells by ∼2.2 FC (Fig. S11C). Next, we measured the *CXCR4* transcript levels using qRT-PCR and observed a significant downregulation by ∼2 to 2.3 FC in c-MET KO-1 and KO-2 cells as compared with control cells (Fig. S11D). We further observed that *CXCR4* mRNA levels were significantly downregulated in primary CD4^+^ T cells and MT-4 cells upon crizotinib treatment (Fig. S11E–F).

Since the transcript levels of *CD4* and *CXCR4* were significantly downregulated following crizotinib treatment and in c-MET KO MT-4 cells, we next examined the cell surface expression of CD4 and CXCR4 proteins upon c-MET KO. We observed that c-MET KO significantly reduced the MFI of CD4-positive cells by ∼2-fold in KO-1 and ∼2.6-fold in KO-2 cells, respectively (Fig. 7D and S12D). We next measured the effect of HGF and crizotinib treatment on the CD4 surface expression levels in primary CD4^+^ T cells and MT-4 cells. We observed no significant change in the CD4 surface expression levels in primary CD4^+^ T cells and MT-4 cells upon HGF treatment (Fig. 7E-F and S12E-F). Consistent with the effects observed in c-MET KO cells, crizotinib treatment also decreased the MFI of CD4-positive cells in primary CD4^+^ T cells (Fig. 7E and S12E) and MT-4 cells (Fig. 7F and S12F) compared with DMSO-treated cells. However, no significant changes were observed in the total percentage of CD4-positive cells following c-MET KO, HGF or crizotinib treatment (Fig. S12A–C).

Further, we measured the surface expression of CXCR4 in control or c-MET KO MT-4 cells. We found that c-MET KO cells exhibited lower MFI of CXCR4-positive cells compared with control MT-4 cells (Fig. 7G and S13D). We next measured the effect of HGF and crizotinib treatment on the CXCR4 surface expression levels in primary CD4^+^ T cells and MT-4 cells. HGF treatment significantly increased the CXCR4 surface expression in primary CD4^+^ T cells or MT-4 cells (Fig. 7H–I and S13E–F). Interestingly, crizotinib treatment reduced the MFI of CXCR4 in primary CD4^+^ T cells (Fig. 7H and S13E) and MT-4 cells (Fig. 7I and S13F). However, no significant change was observed in the total percentage of CXCR4-positive cells upon c-MET KO, HGF or crizotinib treatment (Fig. S13A–C).

### c-MET KO or crizotinib treatment of CD4^+^ T cells decreases receptor-mediated HIV-1 entry

Since the expression of the HIV-1 receptor CD4 and the coreceptor CXCR4 was downregulated by c-MET KO or inhibition in CD4^+^ T cells, we next evaluated HIV-1 entry into cells following c-MET KO or crizotinib treatment. To assess viral entry into cells, control and c-MET KO cells were separately infected with HIV-1 at 37°C for 2 h and cell-associated p24 levels were measured after removing cell-surface attached viruses by extensive washes and trypsin treatment of cells. We observed that c-MET KO significantly impaired HIV-1 entry, with cell-associated p24 levels reduced by ∼2.7-fold and ∼2.4-fold in c-MET KO-1 and KO-2 MT-4 cells respectively, compared with control MT-4 cells (Fig. 7J). Next, we examined the effect of HGF or crizotinib treatment on cell-associated p24 levels in MT-4 cells and in primary CD4⁺ T cells from three independent healthy donors. HGF treatment significantly increased cell-associated p24 levels (∼1.5-fold) in primary CD4⁺ T cells, whereas crizotinib treatment markedly reduced p24 levels by ∼3.5-fold (Fig. 7K). Similarly, HGF-treated MT-4 cells exhibited an approximately 2-fold increase in cell-associated p24 levels, which was reduced by ∼5-fold following crizotinib treatment (Fig. 7L). Collectively, these findings suggest that c-MET disruption significantly reduces HIV-1 entry into CD4⁺ T cells by downregulating viral receptor CD4 and the coreceptor CXCR4.

To further investigate the role of c-MET in post-entry stages of HIV-1 infection, we utilized a single-cycle GFP reporter HIV-1 pseudotyped with vesicular stomatitis virus G protein (VSV-G), which bypasses the requirement for HIV-1-specific receptors and coreceptors during viral entry ^54^. Compared with DMSO-treated control cells, crizotinib treatment did not result in a significant change in GFP expression following infection with VSV-G-pseudotyped HIV-1 in primary CD4^+^ T cells (Fig. S14A–B). Similarly, crizotinib treatment of MT-4 cells did not significantly affect GFP levels following VSV-G-pseudotyped HIV-1 infection compared with DMSO-treated control cells (Fig. S14C–D). Consistent with these findings, c-MET KO cells showed no significant alteration in GFP expression compared with control cells following VSV-G-pseudotyped HIV-1 infection (Fig. S14E–F). These results indicate that c-MET contributes to HIV-1 infection by modulating the expression of key host factors involved in viral entry, including the CD4 receptor and CXCR4 coreceptor.

## Discussion

HIV-1 depends on numerous host cellular and molecular pathways to complete its viral lifecycle. By exploiting host signaling pathways, HIV-1 can fine-tune and rewire these pathways to promote its own replication and survival ^2,6^. HGF/c-MET signaling is well known for its critical role in cancer progression; however, its role in regulating HIV-1 replication in CD4^+^ T cells has not been reported. Our study provides the first evidence demonstrating the importance of HGF/c-MET signaling during HIV-1 infection in primary CD4^+^ T cells (Fig. 8).

**Fig. 8.**
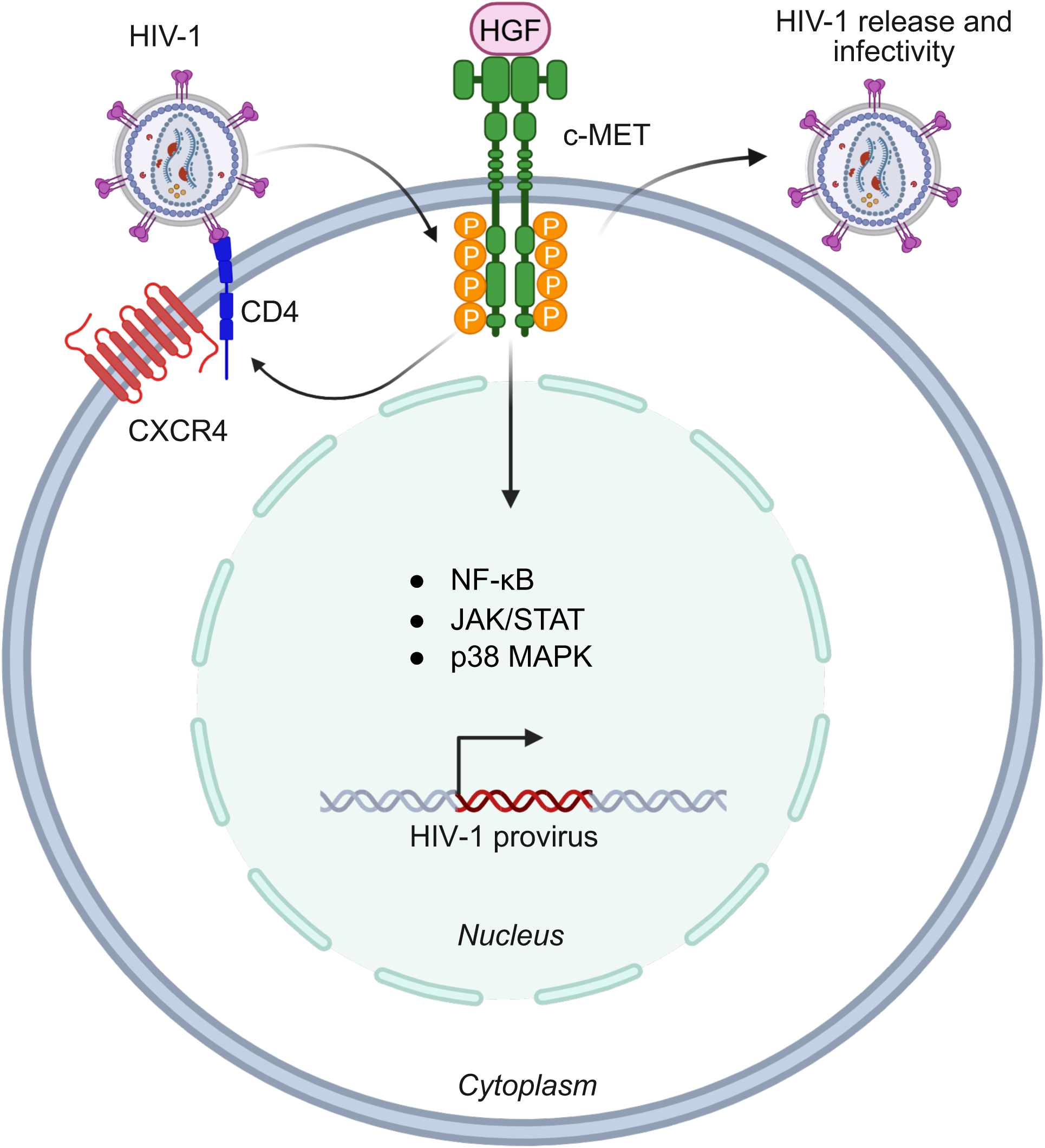
Summary model. Diagram depicts that HIV-1 hijacks the host cell machinery to regulate HGF/c-MET signaling to promote viral replication and cellular responses in CD4^+^ T cells. The figure was created with BioRender.

In our study, we showed that HIV-1 infection in primary CD4^+^ T cells and MT-4 cells induces HGF-mediated upregulation of *c-MET* mRNA expression and enhances c-MET phosphorylation. We demonstrated that HGF treatment significantly increases HIV-1-GFP expression, RT products, Gag mRNA and protein expression, and viral release. However, despite the observed enhancement in viral replication and virion production, HGF treatment resulted in a significant reduction in HIV-1 infectivity. The mechanism by which HIV-1 infection promotes c-MET phosphorylation remains unclear and represents an important area for further investigation. In addition, the precise role of HGF signaling during HIV-1 infection requires deeper exploration, particularly to understand how HGF treatment simultaneously enhances HIV-1 transcription, viral protein expression, while reducing the infectivity of the released virions.

Previous studies have reported that serum HGF levels are elevated during HIV-1 infection in women living with HIV-1 as compared to HIV-1-negative women ^30^. However, to date, no studies have shown the direct effect of HGF treatment on HIV-1 infection. Our study is the first to demonstrate the impact of HGF treatment on HIV-1 replication in primary CD4^+^ T cells. Several earlier studies have suggested an association between elevated HGF levels and disease severity during viral and bacterial infections. For example, increased serum HGF levels have been linked to enhanced liver fibrosis in hepatitis C virus-infected patients ^55^, and these elevated levels do not significantly decline even after successful antiviral therapy ^56^. Moreover, HGF levels are markedly increased in patients with severe influenza A (H1N1) infection, as well as in individuals with inflammatory lung diseases such as interstitial pneumonitis ^57^ and bacterial pneumonia ^58^. Furthermore, patients with COVID-19 have been reported to exhibit significantly elevated circulating HGF levels ^59^, and HGF can predict the disease severity and the mortality.

To further investigate the role of c-MET in HIV-1 replication and infectivity, we utilized the c-MET inhibitor crizotinib. Our findings demonstrated that c-MET inhibition significantly reduced HIV-1 replication, virion production/release, and infectivity. In addition, crizotinib treatment suppressed several HIV-1-induced hosts signaling pathways, including NF-κB, JAK/STAT, and p38 MAPK signaling. Interestingly, Xie et al. reported that the c-MET inhibitor NVP-BVU972 exerts broad-spectrum antiviral activity against several RNA viruses, including vesicular stomatitis virus, encephalomyocarditis virus, and murine hepatitis virus, as well as DNA viruses such as herpes simplex virus type 1 and vaccinia virus ^60^. The antiviral effects of NVP-BVU972 were associated with suppression of NF-κB-mediated inflammatory signaling ^60^. These findings are consistent with our observations demonstrating an antiviral role for crizotinib during HIV-1 infection. Importantly, crizotinib has been approved for cancer treatment in human patients, and several c-MET inhibitors are currently undergoing clinical evaluation for cancer therapy ^61,62^, highlighting the need for further studies to investigate their potential utility during HIV-1 infection and to optimize these inhibitors for antiviral applications.

To further define the mechanistic role of c-MET during HIV-1 infection, we generated c-MET KO MT-4 cells. c-MET KO resulted in a significant reduction in HIV-1-GFP expression, Gag mRNA and protein expression, and p24 release. Notably, virions produced from c-MET KO MT-4 cells exhibited markedly reduced infectivity compared with virions released from control MT-4 cells. To date, relatively few studies have explored the role of HGF/c-MET signaling in viral infections ^63^ and antiviral innate immunity ^64^. Shirasaki et al. demonstrated that leukocyte cell-derived chemotaxin 2, a hepatokine and ligand of c-MET, acts as an antiviral regulator by enhancing innate immune responses and suppressing lymphocytic choriomeningitis virus (LCMV) replication in the liver ^65^. However, this study did not directly examine the role of HGF/c-MET signaling during LCMV infection ^65^. Furthermore, c-MET has also been reported to function as a coreceptor for adeno-associated virus 2 (AAV2) infection by facilitating viral internalization into the cytoplasm ^66^. In another study, c-MET was identified as a novel coreceptor required for AAV3 entry into hepatocytes ^67^. Pharmacological inhibition of c-MET phosphorylation using an inhibitor results in significant reduction of vaccinia virus infection in PM1.CCR5 T cells ^68^. Kaposi sarcoma–associated herpesvirus is known to activate the HGF/c-MET pathway *in vitro* and *in vivo*, and several studies illustrate that c-MET activation is vital for the pathogenesis of Kaposi sarcoma ^69–72^.

HIV-1 infection is known to regulate the NF-κB, JAK/STAT, and p38 MAPK signaling pathways in CD4^+^ T cells ^36–41^. Previous studies have shown that the HGF/c-MET pathway initiates multiple downstream signaling cascades, including the Erk/MAPK, JAK-STAT, and NF-кB signaling pathways and activation of these pathways ^9,10,20,42,43^. Hence, we studied the impact of the HGF/c-MET signaling in regulating these pathways during HIV-1 infection in CD4^+^ T cells. Our result suggested that inhibition of c-MET or c-MET KO significantly reduces the activation of the NF-κB, JAK/STAT, and p38 MAPK signaling pathways. Moreover, our pathway analysis results from the RNA sequencing upon the c-MET KO identified that several pathways related to the inflammatory response, TNF-α signaling through NF-κB, IL2-STAT5, are regulated upon c-MET KO. Previous studies have shown that c-MET plays an important role in regulating the proinflammatory and migratory features of human activated CD4^+^ T cells ^73^. It has been reported that transient pharmacological blockade of c-MET during T cell priming prevents T-cell-mediated inflammation in the heart and led to enhanced survival of heart ^74^. Breville at al. have shown that c-MET is an immune marker of highly pro-inflammatory and migratory CD4^+^ T lymphocytes in both the periphery and central nervous system of multiple sclerosis patients ^75^. Hence, the role of the c-MET in regulating the inflammation during HIV-1 infection in CD4^+^ T cells needs to be investigated.

Our findings provide mechanistic insights into how c-MET KO reduces HIV-1 replication in CD4^+^ T cells (Fig. 8). To identify genes regulated by c-MET, we performed whole-transcriptome sequencing. Notably, transcriptome profiling revealed that *CD4* and *CXCR4* mRNA levels were downregulated upon c-MET KO, which we validated by qRT-PCR. Consistent with the decrease in transcript levels, we observed a significant reduction in CD4 and CXCR4 surface levels following c-MET KO in MT-4 cells. In addition, HGF treatment increased CXCR4 surface expression, whereas treatment with the c-MET inhibitor crizotinib significantly reduced CXCR4 levels. These observations are consistent with previous studies demonstrating that HGF enhances CXCR4 expression and promotes breast cancer invasion and metastasis *in vivo* ^76^. Furthermore, Esencay et al. reported that HGF upregulates CXCR4 expression through NF-κB signaling in glioma cells ^77^. Matteucci et al. also demonstrated differential regulation of CXCR4 by HGF in breast carcinoma cells ^78^. Interestingly, we did not observe a significant change in CD4 surface expression following HGF treatment. However, both crizotinib treatment and c-MET KO resulted in a substantial reduction in CD4 surface levels in MT-4 cells. Collectively, our findings suggest that c-MET signaling plays an important role in regulating CD4 and CXCR4 surface expression in T cells. Nevertheless, the precise mechanisms through which c-MET controls CD4 and CXCR4 transcriptional regulation and surface expression remain to be elucidated. Furthermore, when we used single-cycle HIV-1-pseudotyped with VSV-G protein to infect cells through endocytosis-mediated entry, we did not observe a significant change in HIV-1 infection or gene expression in c-MET KO cells compared with control cells. This result also suggests that HIV-1 receptor-mediated fusion is responsible for the reduced viral infection in c-MET KO cells. Our results indicate that HGF/c-MET signaling promotes HIV-1 replication by positively regulating viral entry.

In summary, we uncovered that the HGF/c-MET signaling pathway plays an important role in regulating the HIV-1 replication and other cell signaling pathways essential for HIV-1 gene expression (Fig. 8). These findings suggest a previously unappreciated function for HGF/c-MET and have significant implications for the regulation of viral and/or host cellular proteins in HIV-1-infected cells. Thus, the HGF/c-MET signaling pathway emerged as a critical axis to regulate the HIV-1 infection via modulating CD4 and CXCR4 levels.

## Materials and Methods

### Ethics statement

The Institutional Review Board (IRB) at the University of Iowa has approved the *in vitro* experiments in this study involving human blood cells from de-identified healthy donors. The consent requirements for the de-identified blood samples were waived by IRB.

### Cell culture

MT-4 and primary CD4^+^ T cells were cultured in RPMI-1640 (ATCC) supplemented with 10% fetal bovine serum (FBS; R&D Systems) and antibiotics (100 U/mL penicillin and 100 μg/mL streptomycin, Gibco). HEK293T, and TZM-bl cells were cultured in DMEM (Gibco) with 10% FBS and antibiotics as described ^79^. All cells were cultured at 37°C with 5% CO2 and tested negative for mycoplasma contamination using a PCR-based universal mycoplasma detection kit (ATCC 30-1012K). Healthy deidentified donor blood was purchased from the DeGowin Blood Center at the University of Iowa. PBMCs were isolated from healthy donor blood using Ficoll-Paque PLUS (17144002, Cytiva). CD4^+^ T cells were enriched using EasySep Human CD4^+^ T cell isolation kit (17952, STEMCELL Technologies) and activated using ImmunoCult Human CD3/CD28/CD2 T cell activator (10970, STEMCELL Technologies) for 72 h as described ^79^. The recombinant human HGF (PHG0254, Gibco) was used to treat CD4^+^ T cells with a concentration of 10 ng/mL and was replenished every 12 h. Cells were pre-treated with crizotinib (or PF-2341066) at a concentration of 0.4 µM (S1068, Selleckchem) before mock or HIV-1 infection or DMSO vehicle control.

### HIV-1 production and infection

Replication-competent X4-tropic HIV-1, p-NLENG-IRES HIV-1 were generated by transfection of HEK293T cells using PolyFect Transfection Reagent (301105) as described ^80^. The supernatants were filtered (0.45 μm) and digested with DNase I (60 U/mL, Turbo, Invitrogen) for 30 min at 37°C. The viral stock titters were calculated through serial dilution on MT-4 cell lines. For HIV-1 infection, MT-4 and primary activated CD4^+^ T cells were infected with HIV-1 at a multiplicity of infection (MOI) of 1. Cells were washed twice with Dulbecco’s phosphate-buffered saline (DPBS) and resuspended in a fresh culture medium. To inhibit HIV-1 infection, cells were treated with the fusion inhibitor AMD3100 (10 μM) and reverse transcriptase inhibitor NVP (10 μM). Both inhibitors were obtained from the AIDS Research and Reference Reagent Program, NIH. HIV-1 supernatant p24 levels were detected by p24 ELISA using anti-p24-coated plates (AIDS and Cancer Virus Program, National Cancer Institute, Frederick, MD) as described ^81^. Single-cycle GFP reporter HIV-1 pseudotyped with VSV-G was generated by transfection of HEK293T cells as described ^54^.

### Antibodies and immunoblotting

Antibodies used for immunoblotting were as follows: HIV-1 p24 (clone #24-2, the AIDS Research and Reference Reagent Program, NIH), c-MET (8198, Cell Signaling), phospho-Met-Tyr1234/1235 (3077, Cell Signaling), phospho-Met-Tyr1349 (3121, Cell Signaling), phospho-c-Met (Tyr1356) (PA5-40218, Invitrogen), Tubulin (ab7291, Abcam), IKKα (2682, Cell Signaling), IKKα (61294S, Cell Signaling), IKKβ (8943, Cell Signaling), phospho-IKKα/β (Ser176/180) (2697, Cell Signaling), IκBα (4814, Cell Signaling), phospho-IκBα (Ser32/36) (9246, Cell Signaling), phospho-p38 (4511T, Cell Signaling), p38 MAPK (8690T, Cell Signaling), phospho-STAT1 (8826, Cell Signaling), STAT1 (9172, Cell Signaling), phospho-STAT3 (9134, Cell Signaling), and STAT3 (9139, Cell Signaling). Secondary antibodies used were goat anti-mouse IgG (H + L) HRP (W402B, Promega) and goat antirabbit lgG (H + L) HRP (W401B, Promega). Cells were harvested and lysed in cell lysis buffer (9803, Cell Signaling Technology) with a protease and phosphatase inhibitor (A32959, Pierce, Thermo Scientific). Immunoblotting was performed as described ^82^. Tubulin was used as a loading control for all immunoblots.

### Quantitative PCR (qPCR) assays and primer sequences

Total RNA was extracted using TRIzol (Invitrogen) or RNeasy Plus Kit (74134, Qiagen). cDNA was synthesized from the extracted RNA using iScript cDNA Synthesis Kit (1708891, Bio-Rad), and qPCR was performed using iTaq Universal SYBR Green Supermix (1725120, Bio-Rad). Sequences of qRT-PCR primers (Integrated DNA Technologies) are listed as follows.

CD4 forward primer (FP): CCTCCTGCTTTTCATTGGGCTAG

CD4 reverse primer (RP): TGAGGACACTGGCAGGTCTTCT

CXCR4 FP: CTCCTCTTTGTCATCACGCTTCC

CXCR4 RP: GGATGAGGACACTGCTGTAGAG

c-MET FP: TGCACAGTTGGTCCTGCCATGA

c-MET RP: CAGCCATAGGACCGTATTTCGG

HPRT FP: CATTATGCTGAGGATTTGGAAAGG

HPRT RP: CTTGAGCACACAGAGGGCTACA

To quantify HIV-1 early and late RT products after viral infection, cellular DNA was extracted using a QIAamp DNA Blood Mini kit (Qiagen) and 50 ng DNA was used as a template for qPCR. Because HIV-1 reverse transcription initiates from PBS, early RT products were quantified using primers ert2f (5-GTGCCCGTCTGTTGTGTGAC-3) and ert2r (5-GGCGCCACTGCTAGAGATTT-3), which amplify a fragment overlapping with the HIV-1 primer-binding site and R regions with 74 and 7 nt, respectively. The late RT products were amplified with an LTR R-specific primer (forward, 5-GGGAGCTCTCTGGCTAACT-3) and a gag-specific primer (reverse, 5-GGATTAACTGCGAATCGTTC-3) as described ^83^. Unspliced GAPDH levels were also quantified to normalize early and late RT data ^83^.

### c-MET KD in activated primary CD4^+^ T cells

Activated primary CD4^+^ T cells were nucleofected with CRISPR-Cas9 ribonucleoprotein (crRNP) as described ^79,84^. Synthetic crRNA, trans-activating crRNA (tracrRNA), and recombinant Cas9 protein were purchased from Integrated DNA Technologies. Briefly, activated primary CD4^+^ T cells were resuspended in P3 buffer (Lonza) at a density of 5 × 10^5^ cells per reaction and mixed with the crRNP complex. Electroporation was performed using the 4D-Nucleofector System (Lonza) with program EH115. After nucleofection, cells were transferred to pre-warmed RPMI-1640 medium containing 20% FBS and incubated at 37°C. Knockout efficiency was evaluated 72 h post-electroporation by immunoblotting. The sequences of negative control and *c-MET* crRNA are listed below:

Negative control crRNA: CGTTAATCGCGTATAATACG

cr-C-MET-(KD-1): TTACTTCTTGACGGTCCAAA

cr-C-MET-(KD-2): TCAGCTTCCCAACTTCACCG

### Generation of stable MT-4 cell lines with c-MET KO

The *c-MET* sgRNA (KO-1: TCAGCTTCCCAACTTCACCG, KO-2: TGCATTCGATATCAGTGAGA) and control sgRNA CACCGCGCTTCCGCGCCCGTTCAA were purchased from Applied Biological Materials (LG128379). MT-4 cells were transduced with lentiviruses in the presence of polybrene (10 μg/mL) by spinoculation at 1,200 × *g* for 2 h at 25°C. Transduced cells were cultured in complete RPMI-1640 for 48 h prior to selection with puromycin (1 μg/mL). After 7 days of selection, single-cell clones were obtained by limiting dilution. c-MET KO MT-4 cells (KO-1 and KO-2 clones) were confirmed by immunoblotting.

### HIV-1 infectivity measurement in TZM-bl cells

TZM-bl cells (1 × 10^5^) were seeded in 24-well plates overnight and infected with 2 ng p24 HIV-1 stocks. After 48 h, the luciferase activity was measured using ONE-Glo EX Luciferase Assay System (E8120, Promega). Luminescence was quantified using a microplate reader and normalized to total protein content.

### RNA sequencing

Quality of the fastq files was assessed using FastQC v0.12.1. Reads were then quality filtered using fastp v0.24.0 with poly-X tail trimming, 3’ quality-based tail trimming, a minimum Phred quality score of 15, and a minimum length requirement of 50 bp. Quality-filtered reads were aligned to the reference genome using STAR aligner v2.7.11 with non-canonical splice junction removal and output of unmapped reads, followed by coordinate sorting using sam tools v1.22.1. PCR and optical duplicates were removed using UMI-based deduplication with UMIcollapsev1.1.0. Alignment quality metrics, strand specificity, and read distribution across genomic features were assessed using RSeQC v5.0.4 and Quali map v2.3, with results aggregated into a comprehensive quality control report using MultiQCv1.32. Gene-level expression quantification was performed using feature Counts (subread package v2.1.1) with strand-specific counting, multi-mapping read fractional assignment, exons and three prime UTR as the feature identifiers, and grouped by gene_id. Final gene counts were annotated with gene biotype and other metadata extracted from the reference GTF file. Sample-sample correlations for sample-sample heatmap and PCA were calculated on normalized counts (TMM, trimmed mean of M-values) using Pearson correlation. Differential expression was done with edgeR v4.0.16 using standard practice including filtering for low-expressed genes with edgeR filterByExpr with default values. Functional enrichment, when available, is performed using gene set enrichment analysis with gseapy v0.12 using the MSigDB Hallmark gene set.

### HIV-1 entry assay

To measure HIV-1 entry into cells, primary CD4^+^ T cells and MT-4 cells were infected with HIV-p-NLENG-IRES HIV-1 for 2 h at 37°C. Following infection, cells were washed with 1x DPBS, to test the protease sensitivity of cell-associated HIV-1, infected cells were treated with 0.25% trypsin (Invitrogen) at room temperature for 4 min, and cells were subsequently neutralized with culture medium containing 10% FBS and washed prior to cell lysis. Cells were then lysed with 200 μl of 1% Triton X-100 buffer for Gag p24 quantification by ELISA as previously described ^54^.

### Flow cytometry and antibodies

To assess cell surface expression of CD4 and CXCR4, primary CD4⁺ T cells and MT-4 cells (1 × 10O cells) were first washed and stained with live dead stain using Ghost Dye Violet 450 (SKU 13-0863-T500 for 30 minutes at 4 °C, followed by washing with FACS buffer. Cells were then fixed with 4% PFA for 30 minutes at 37 °C and washed again with FACS buffer. Subsequently, cells were stained with APC Mouse Anti-Human CD4 (BD Pharmingen, 555349) or APC-conjugated anti-human CD184 (CXCR4) antibody (Miltenyi Biotec, 130-123-814), followed by three washes with FACS buffer and resuspended in DPBS. Stained cells were analyzed using a Beckman CytoFLEX cytometer.

### Statistical analysis

Data were analyzed using Paired T-test and analysis of variance (ANOVA) with Prism software and statistical significance was defined as * *P* <0.05, ** *P* <0.01, *** *P* <0.001, **** *P* <0.0001.

## Acknowledgments

We thank the Wu lab members for helpful discussions and suggestions. We appreciate the reagents provided by the National Institutes of Health (NIH) AIDS Reagent Program.

## Funding

This work was supported by NIH grants R33AI169659 and P30CA086862-25S (to L.W.). L.W. and his lab are also supported by NIH grants R01AI189220 and R21AI181742. The content is solely the responsibility of the authors and does not necessarily represent the official views of the NIH.

## Author contributions

S.T. and L.W. designed and conceptualized the study. S.T. performed all experiments and analyzed data. M.M.A. provided technical support for the experiments. S.T. drafted the original manuscript and L.W. edited the manuscript. L.W. supervised the study and acquired funding.

## Competing Interests

The authors (S.T. and L.W.) have filed a provision patent application (64/069,456) on May 19, 2026 regarding the methods of inhibiting HIV-1 replication that were described in this work.

## Data availability

The RNA sequencing data were deposited in the Gene Expression Omnibus (GEO) with an accession number GSE335060 and the release date of September 1, 2026. All other data needed to evaluate the conclusions in the paper are present in the paper or the Supplementary Materials.

## Supplementary figure legends

**Fig. S1.**
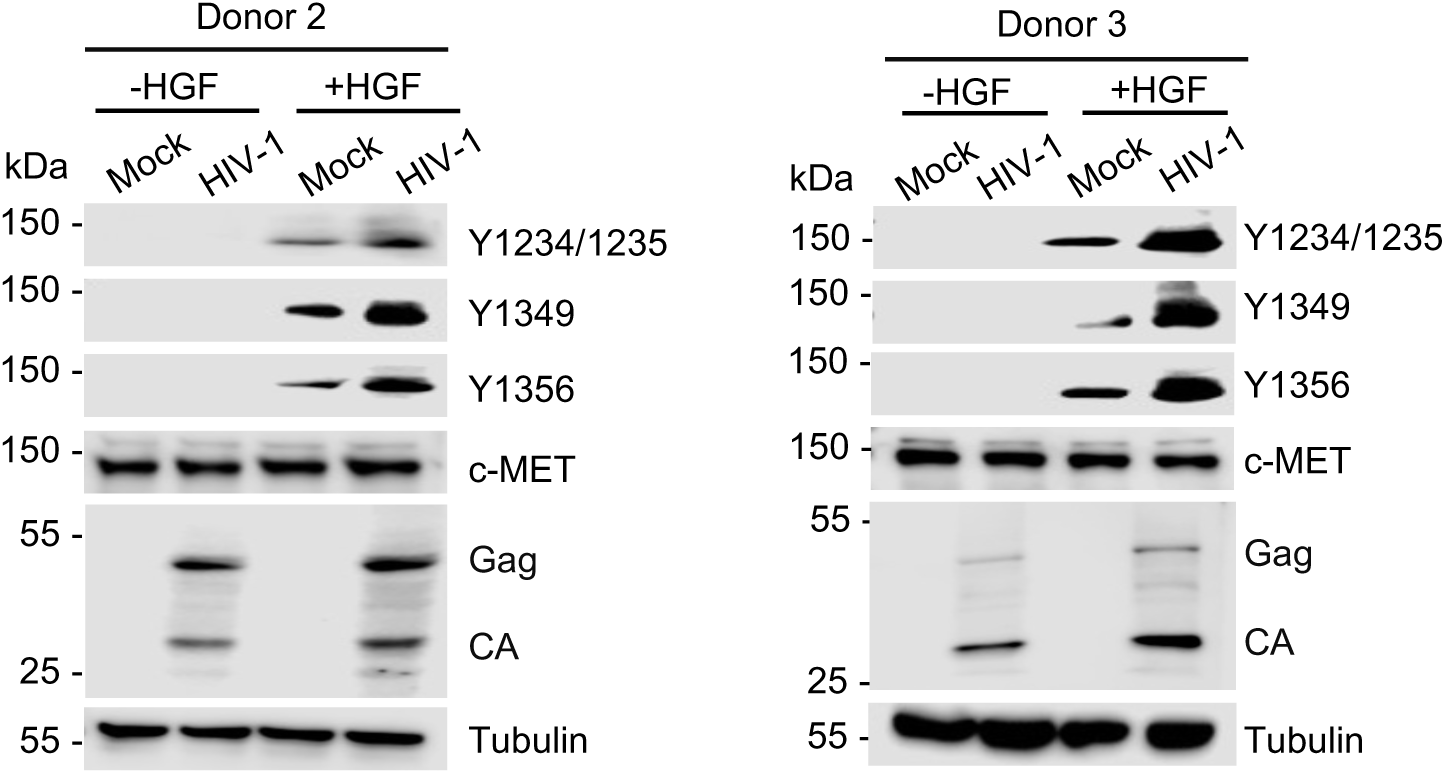
HIV-1 infection promotes HGF-mediated phosphorylation of c-MET in primary CD4^+^ T cells from donors 2 and 3. Detection of phosphorylated Y1234/1234, Y1349, and Y1356, c-MET, HIV-1 Gag and CA, and tubulin by immunoblotting. Tubulin was used as a loading control.

**Fig. S2.**
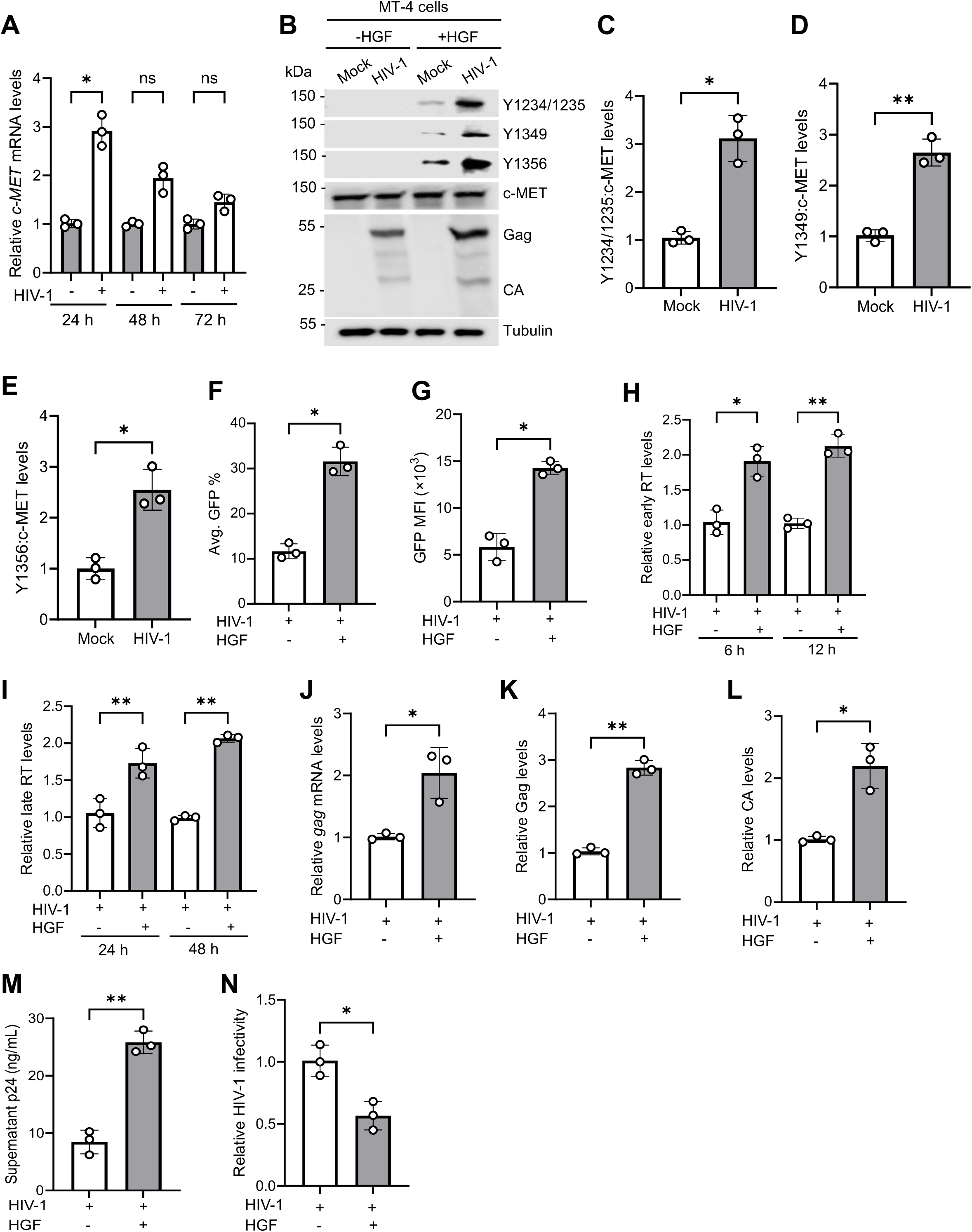
HIV-1 infection promotes HGF-mediated phosphorylation of c-MET in MT-4 cells. **(A)** Time course analysis of the *c-MET* mRNA in MT4 cells at indicated time points during HIV-1 infection. **(B)** Detection of phosphorylated Y1234/1234, Y1349, and Y1356, c-MET, HIV-1 Gag and CA, and tubulin by immunoblotting. Tubulin was used as a loading control. **(C-E)** The relative levels of phosphorylated Y1234/1235 (C), Y1349 (D), Y1356 (E) were quantified by densitometry analysis and normalized to tubulin. The level of proteins in mock-infected HGF treated cells was set to 1. **(F-G)** HIV-1 infection was measured by GFP expression using flow cytometry at 48 hpi. Changes in average GFP percentage (F) and change in MFI of GFP-positive cells (G) from HIV-1 infected MT4 cells with or without HGF treatment are shown. **(H-I)** HIV-1 early reverse transcription (early RT) products (H) and late RT products (I) were measured by qPCR at the indicated time points in HIV-1 infected MT-4 cells treated with or without HGF. Unspliced *GAPDH* was used for normalization. **(J)** Total RNA was isolated from HIV-1 infected MT-4 cells with or without HGF treatment 48 hpi, and HIV-1 *gag* mRNA levels were quantified using qRT-PCR. The amplification of *HPRT* was used as an internal control. **(K-L)** The Gag (K) and CA (L) levels were quantified by densitometry analysis and normalized to tubulin. The level of Gag and CA proteins from HIV-1-infected MT-4 cells was set to 1. **(M)** Cell culture supernatants were collected from HIV-1 infected MT-4 cells with or without HGF treatment, and p24 levels were quantified by ELISA. **(N)** TZM-bl cells were infected with the supernatant of HIV-1 infected MT-4 cells with or without HGF treatment, and luciferase activity was measured at 48 hpi. Paired T-test and one-way ANOVA multiple comparisons test were used to evaluate the statistical significance of the difference between sample groups. * *P* <0.05, ** *P* <0.01, ns, not significant.

**Fig. S3.**
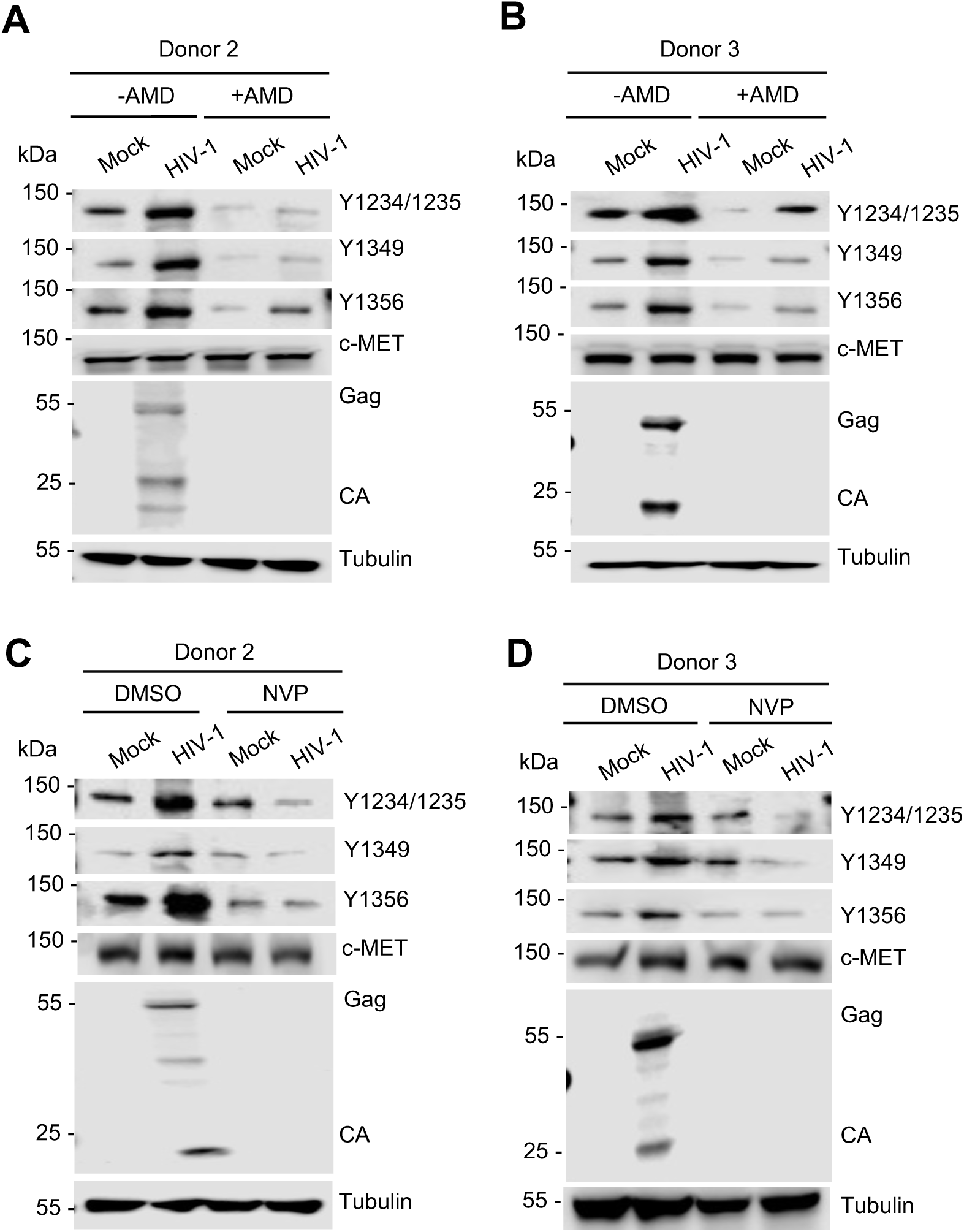
AMD or NVP treatment decreases phosphorylation of c-MET levels in HGF-treated primary CD4^+^ T cells (donors 2 and 3) during HIV-1 infection. **(A-D)**. Detection of phosphorylated Y1234/1235, Y1349, and Y1356 of c-MET, total c-MET, HIV-1 Gag, CA, and tubulin by immunoblotting in mock or HIV-1-infected primary CD4^+^ T cells treated with DMSO or AMD **(A-B)** or NVP **(C-D)**. Tubulin was used as a loading control.

**Fig. S4.**
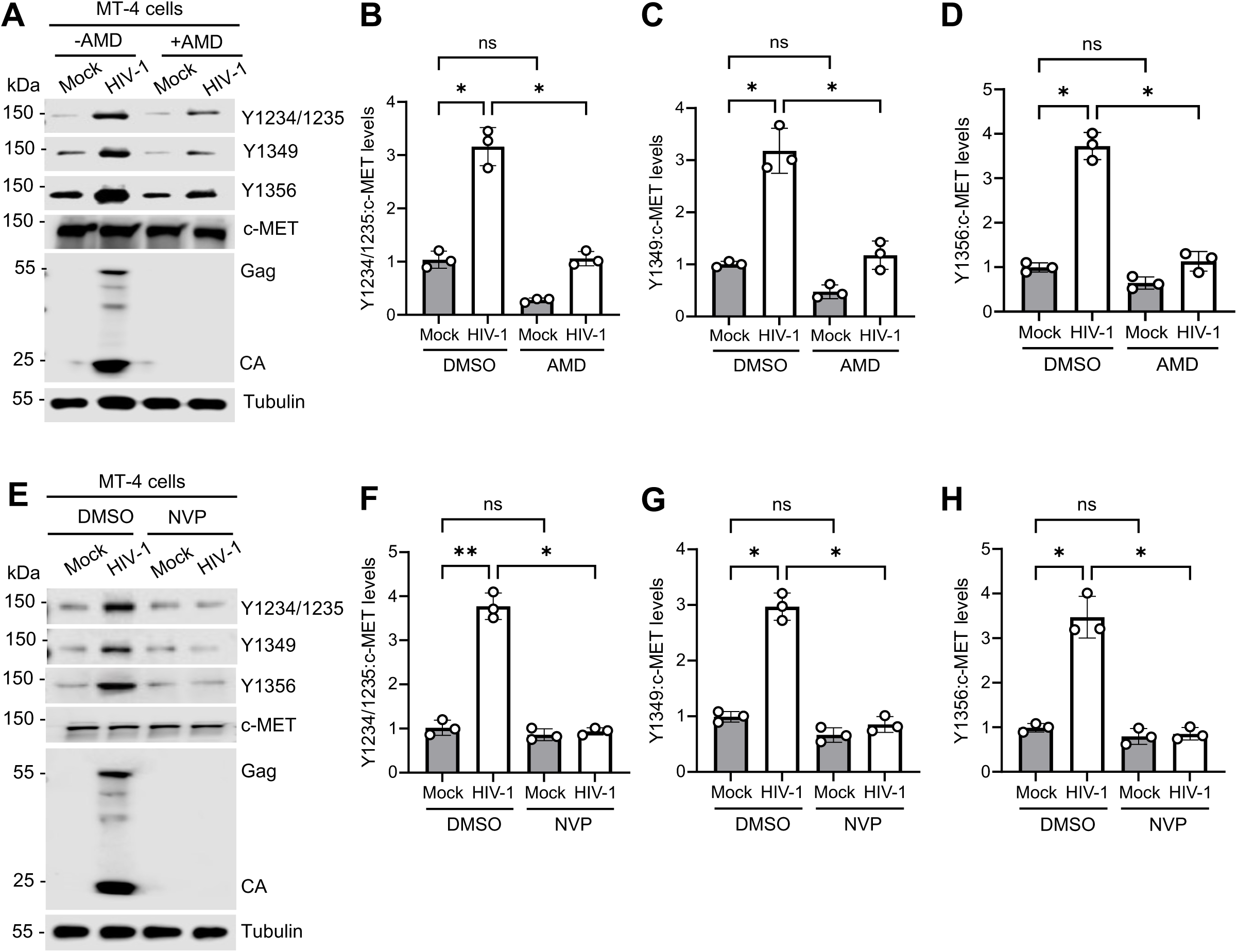
AMD or NVP treatment decreases phosphorylation of c-MET levels in HGF treated MT-4 cells during HIV-1 infection. **(A** and **E)** Detection of phosphorylated Y1234/1235, Y1349, and Y1356 of c-MET, total c-MET, HIV-1 Gag, CA, and tubulin by immunoblotting in mock or HIV-1 infected MT-4 cells treated with DMSO, AMD (A) or NVP (E). Tubulin was used as a loading control. **(B-D and F-H)** The relative levels of phosphorylated Y1234/1235 (B and F), Y1349 (C and G), Y1356 (D and H) of AMD (B-D) or NVP (F-H) treated cells were quantified by densitometry analysis and normalized to total c-MET and tubulin. The level of proteins in mock-infected DMSO treated cells was set to 1. Two-way ANOVA, Šídák’s multiple comparisons test was used to evaluate the statistical significance of the difference between sample groups. * *P* <0.05, ** *P* <0.01, ns, not significant.

**Fig. S5.**
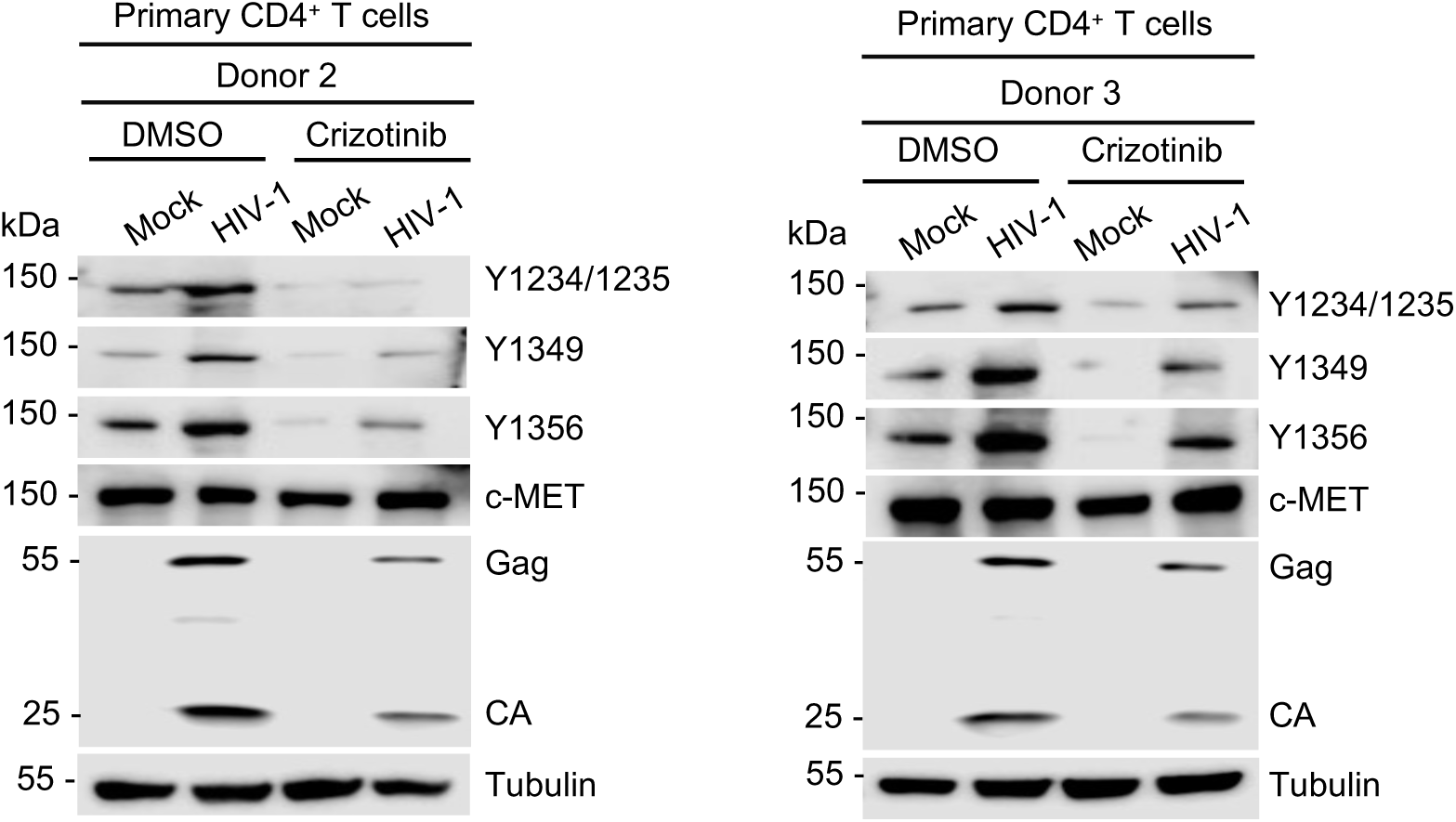
Crizotinib treatment decreases HIV-1 replication in HGF-treated primary CD4^+^ T cells from donors 2 and 3. Detection of phosphorylated Y1234/1234, Y1349, and Y1356, c-MET, HIV-1 Gag, CA, and tubulin by immunoblotting in mock or HIV-1 infected primary CD4^+^ T cells treated with DMSO or crizotinib. Tubulin was used as a loading control.

**Fig. S6.**
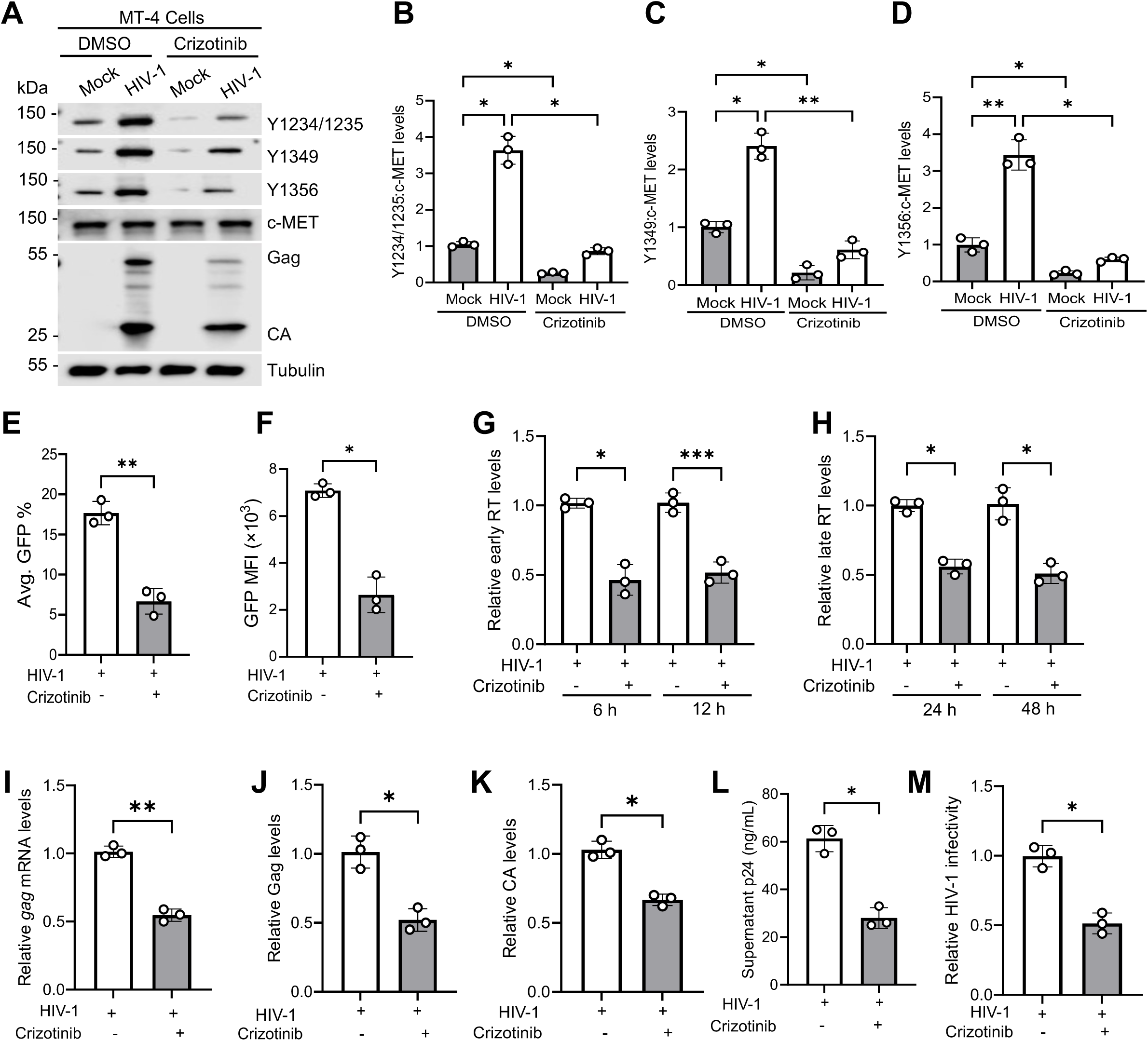
Crizotinib treatment decreases HIV-1 replication and infectivity in HGF-treated MT-4 cells. **(A)** Detection of phosphorylated Y1234/1234, Y1349, and Y1356, c-MET, HIV-1 Gag, CA, and tubulin by immunoblotting in mock or HIV-1 infected MT-4 cells treated with DMSO or crizotinib. Tubulin was used as a loading control. **(B-D)** The relative levels of phosphorylated Y1234/1235 (B), Y1349 (C), Y1356 (D) were quantified by densitometry analysis and normalized to tubulin. The level of proteins in mock-infected HGF treated cells was set to 1. **(E-F)** HIV-1 infection was measured by GFP expression using flow cytometry at 48 hpi. Changes in average GFP percentage (E) and MFI of GFP-positive cells (F) from HIV-1 MT-4 cells treated with DMSO or crizotinib. **(G-H)** HIV-1 early RT products **(G)** and late RT products **(H)** were measured by qPCR at the indicated time points in HIV-1 infected MT-4 cells treated with DMSO or crizotinib. Unspliced *GAPDH* was used for normalization. **(I)** Total RNA was isolated from HIV-1 infected MT-4 cells treated with DMSO or crizotinib, and HIV-1 *gag* mRNA levels were quantified using qRT-PCR. The amplification of *HPRT* was used as an internal control for RT-PCR. **(J-K)** The Gag **(J)** and CA **(K)** levels were quantified by densitometry analysis and normalized to tubulin. The level of Gag and CA proteins from mock DMSO treated cells was set to 1. **(L)** Cell culture supernatants were collected from HIV-1 infected MT-4 cells with treated with DMSO or crizotinib, and p24 levels were quantified by ELISA. **(M)** TZM-bl cells were infected with the supernatant of HIV-1 infected MT-4 cells treated with DMSO or crizotinib, and HIV-1 infectivity was measured by luciferase activities at 48 hpi. Paired T-test and one-way ANOVA multiple comparisons test were used to evaluate the statistical significance of the difference between sample groups. * *P* <0.05, ** *P* <0.01, *** *P* <0.001.

**Fig. S7.**
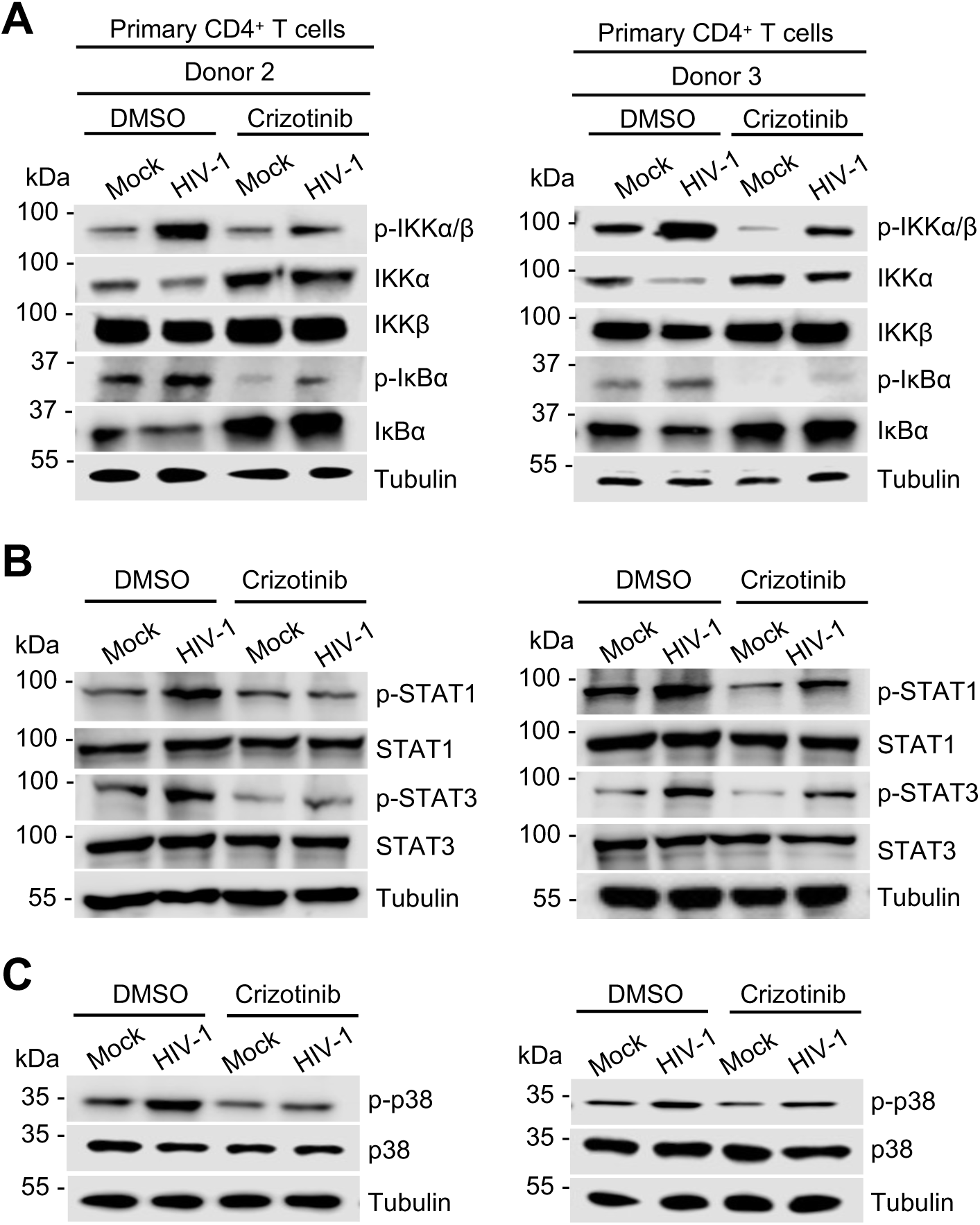
Crizotinib treatment reduces p-IKKα/β, IκBα, p-STAT1, p-STAT3, and p-p38 MAPK in HGF-treated primary CD4^+^ T cells (donors 2 and 3) during HIV-1 infection. **(A-C)** Mock or HIV-1 infected primary CD4^+^ cells were treated with DMSO or crizotinib. The cell lysates were harvested and **(A)** p-IKKα/β, IKKα, IKKβ, p-IκBα, IκBα, tubulin **(B)** p-STAT1, STAT1, p-STAT3, STAT3, tubulin, and **(C)** p-p38, p38, and tubulin were detected by immunoblotting. Tubulin was a loading control.

**Fig. S8.**
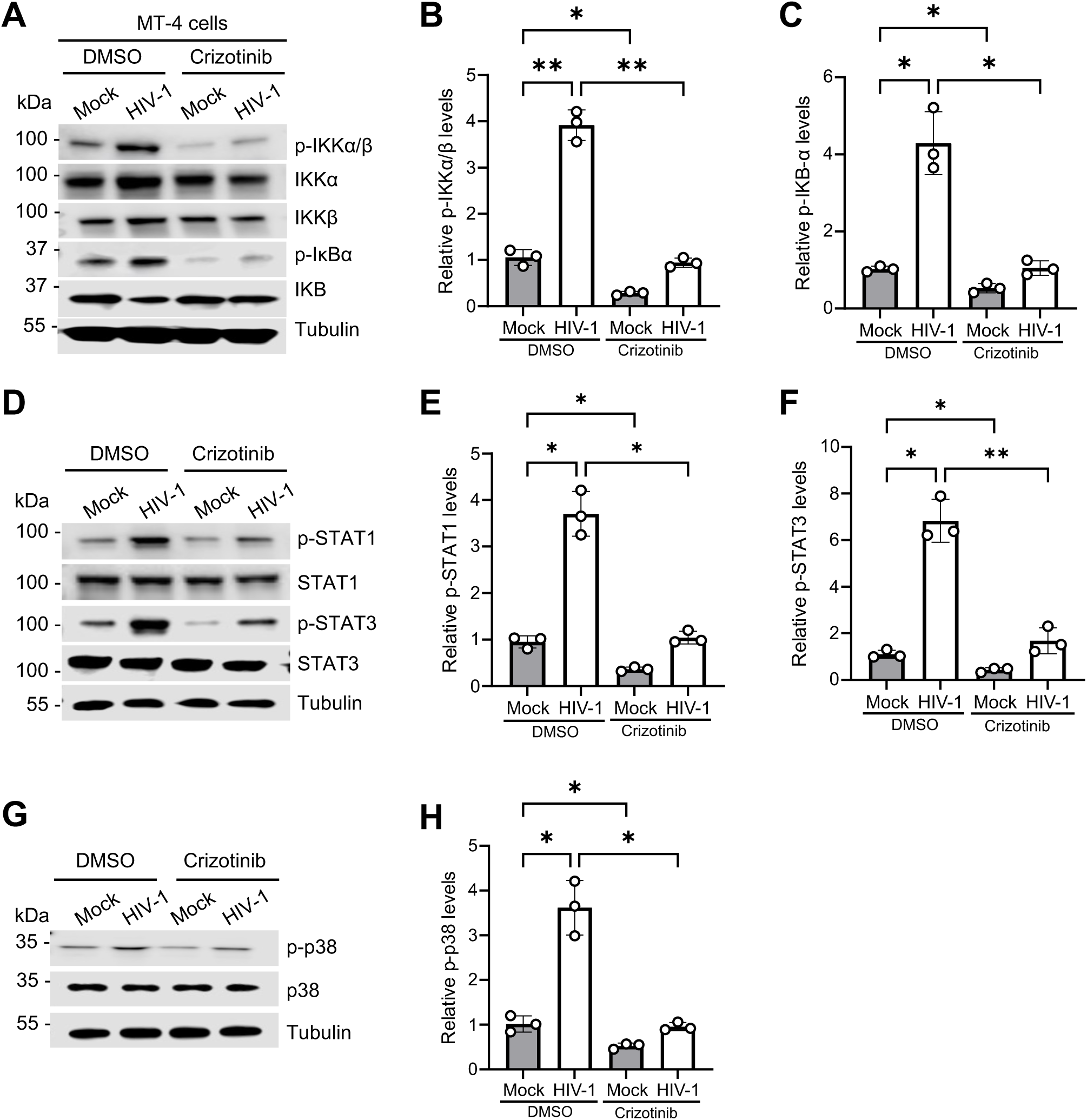
Crizotinib treatment reduces p-IKKα/β, IκBα, p-STAT1, p-STAT3, and p-p38 MAPK in HGF-treated MT-4 cells during HIV-1 infection. **(A, D, and G)** Mock or HIV-1 infected MT-4 cells were treated with DMSO or crizotinib. The cell lysates were harvested and (A) p-IKKα/β, IKKα, IKKβ, p-IκBα, IκBα, and tubulin (D) p-STAT1, STAT1, p-STAT3, STAT3, tubulin and (G) p-p38, p38, and tubulin were detected by immunoblotting. Tubulin was a loading control. The relative p-IKKα/β **(B)**, p-IκBα **(C)**, p-STAT1 **(E)** p-STAT3 **(F)**, and p-p38 **(H)** levels were quantified by densitometry analysis. Relative levels were normalized to tubulin. The level of proteins in mock-infected DMSO treated cells was set to 1. Two-way ANOVA, Šídák’s multiple comparisons test was used to evaluate the statistical significance of the difference between sample groups. * *P* <0.05, ** *P* <0.01.

**Fig. S9.**
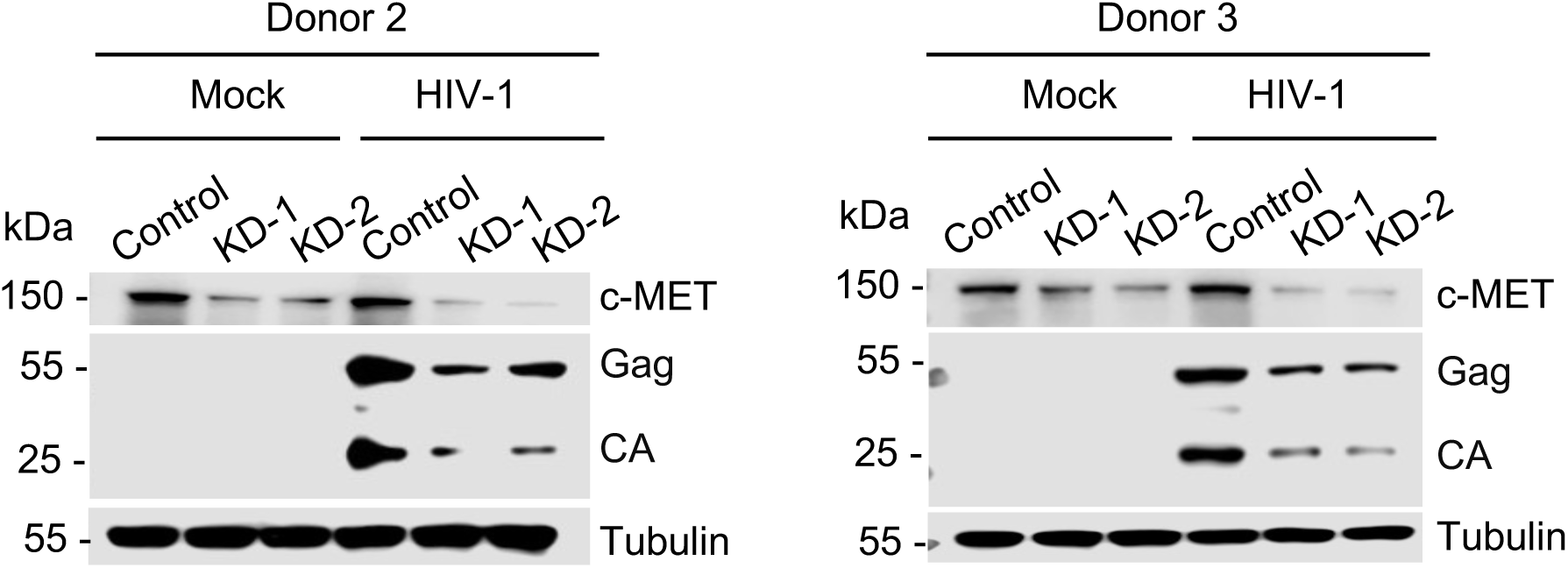
c-MET KD in primary CD4^+^ T cells reduces c-MET, HIV-1 Gag and CA protein levels. KD of c-MET expression in activated primary CD4^+^ T cells from three donors using crRNP targeting two different sites in *c-MET* (KD-1 and KD-2). The cell lysates were harvested from donor 2 and 3 and the expressions of c-MET, HIV-1 Gag and CA were detected by immunoblotting. Tubulin was a loading control.

**Fig. S10.**
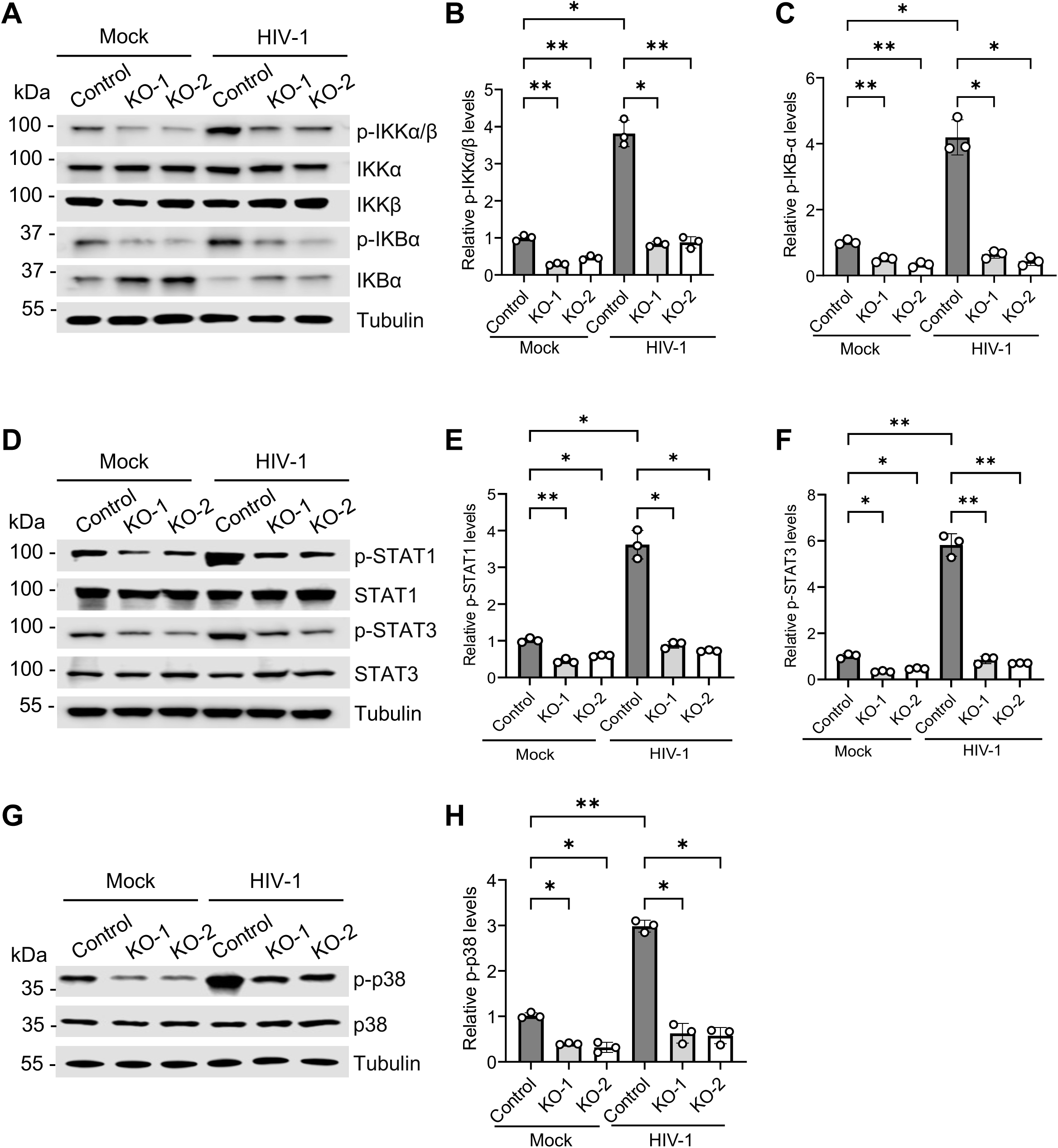
c-MET KO in MT-4 cells decreases p-IKKα/β, IκBα, p-STAT1, p-STAT3, and p-p38 MAPK during HIV-1 infection. **(A, D and G)** MT-4 control and c-MET KO cells were mock or HIV-1 infected. The cell lysates were harvested and (A) p-IKKα/β, IKKα, IKKβ, p-IκBα, IκBα, and tubulin, (D) p-STAT1, STAT1, p-STAT3, STAT3 and tubulin, (G) p-p38, p-38, and tubulin were detected by immunoblotting. Tubulin was a loading control. Results from one of three independent experiments are shown. **(B, C, E, F** and **H)** Quantification results of three independent experiments. The relative p-IKKα/β (B), p-IκBα (C), p-STAT1 (E), p-STAT3 (F), p-p38 (H) levels were quantified by densitometry analysis. Relative levels were normalized to the tubulin. Two-way ANOVA, Šídák’s multiple comparisons test was used to evaluate the statistical significance of the difference between sample groups. * *P* <0.05, ** *P* <0.01.

**Fig. S11.**
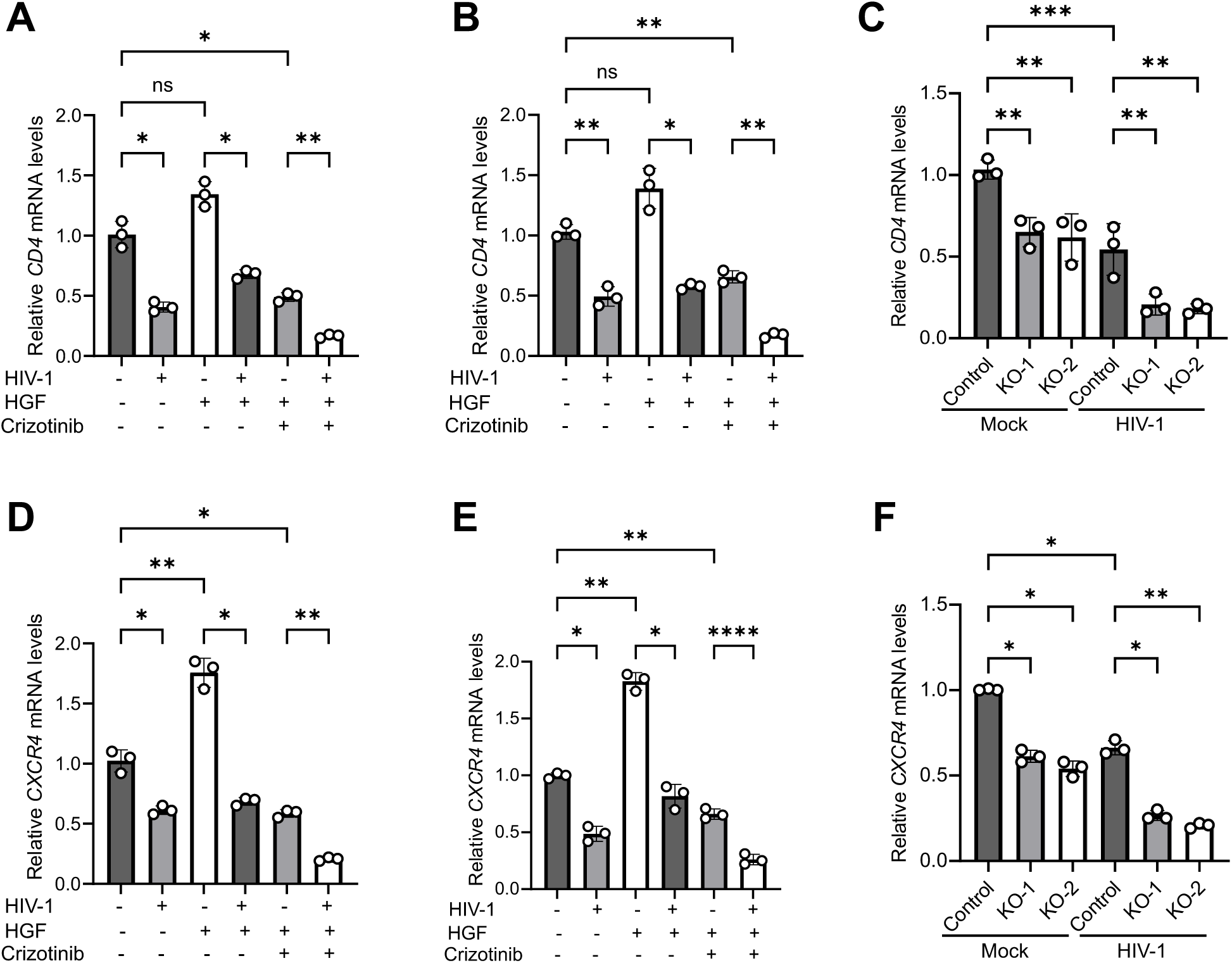
Crizotinib treatment or c-MET KO reduces *CD4* and *CXCR4* transcript levels in CD4^+^ T cells. **(A-D)** Total RNA was isolated from mock, or HIV-1 infected three independent donor primary CD4^+^ T cells (A-B) MT-4 cells (C-D) treated either with HGF, DMSO or crizotinib, *CD4* mRNA levels (A and C) and *CXCR4* mRNA levels (B and D) were quantified using qRT-PCR. HPRT was used as an internal control. **(E-F)** Total RNA was isolated from mock or HIV-1 infected control or c-MET KO MT-4 cells, *CD4* mRNA (E) and *CXCR4* mRNA (F) levels were quantified using qRT-PCR. HPRT was used as an internal control. Two-way ANOVA, Šídák’s multiple comparisons test was used to evaluate the statistical significance of the difference between sample groups. * *P* <0.05, ** *P* <0.01, *** *P* <0.001, **** *P* <0.0001, ns, not significant.

**Fig. S12.**
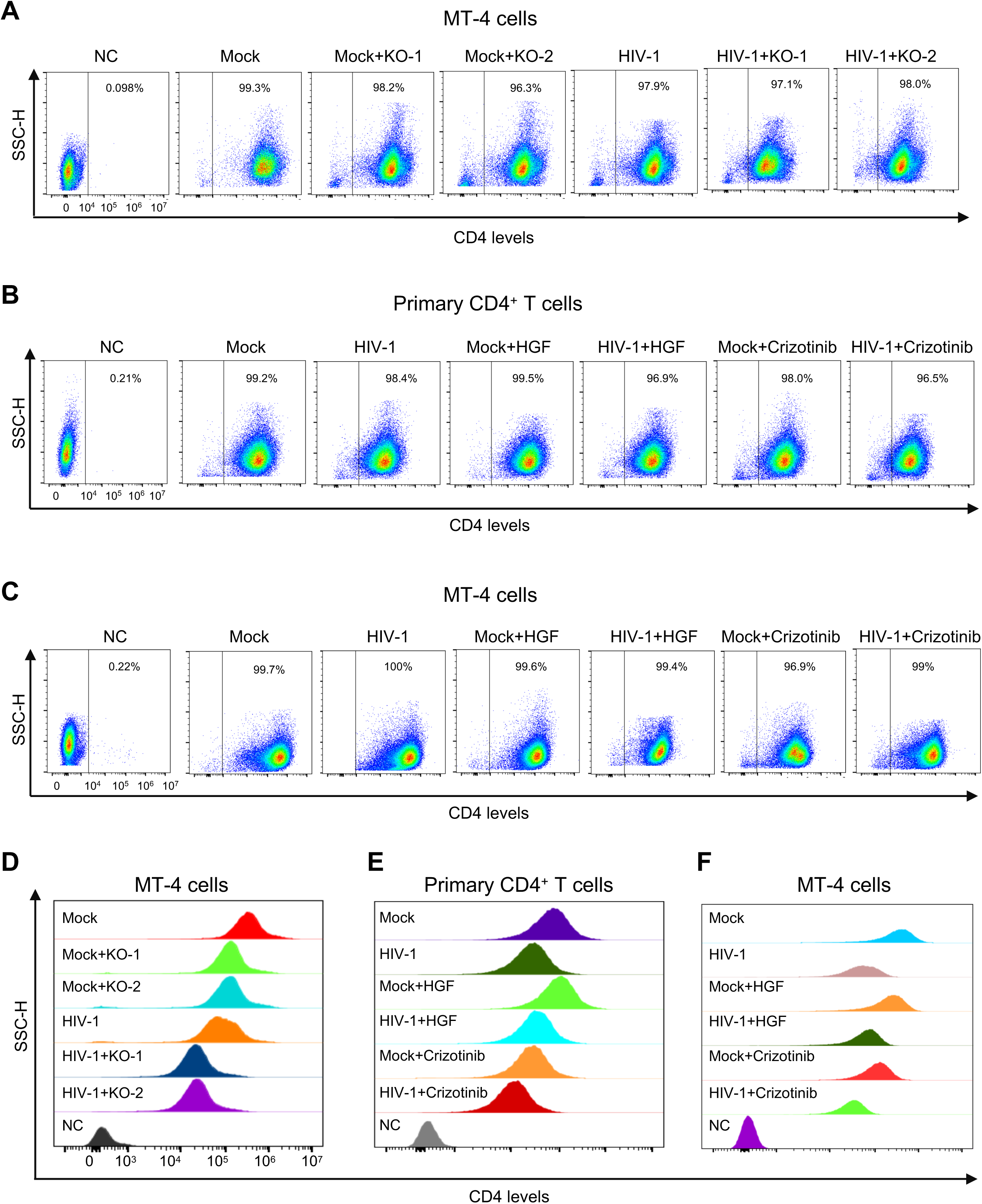
Crizotinib treatment or c-MET KO reduces CD4 surface expression in CD4^+^ T cells. **(A-C)** The cell surface expression of CD4 was measured by flow cytometry in mock or HIV-1 infected control and c-MET KO cells (A) mock or HIV-1-infected primary CD4^+^ T cells (B) or MT-4 cells (C) treated with HGF or crizotinib. The percentage of CD4 positive cells are shown. **(D-F)** The histogram peaks of CD4 staining levels on control or c-MET KO MT-4 cells (D), primary CD4^+^ T cells (E), and MT-4 cells (F) are shown. The results are representative of three independent experiments performed with biological samples (n=3) or primary CD4^+^ T cells from 3 independent donors.

**Fig. S13.**
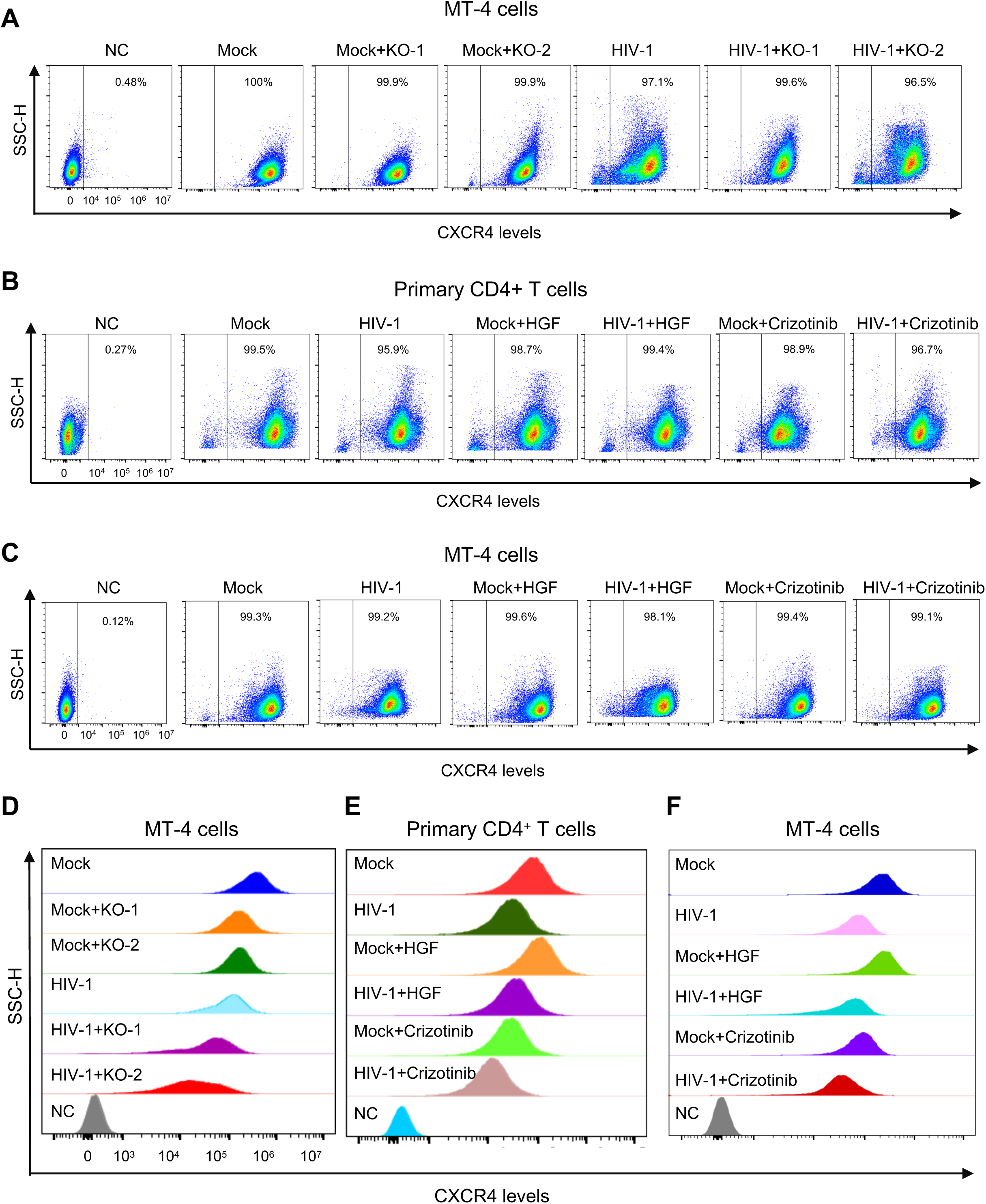
Crizotinib treatment or c-MET KO reduces CXCR4 surface expression in CD4^+^ T cells. **(A-C)** The cell surface expression of CXCR4 was measured by flow cytometry in mock or HIV-1 infected control and c-MET KO cells (A) mock or HIV-1-infected primary CD4^+^ T cells (B) or MT-4 cells (C) treated with HGF or crizotinib. The percentage of CXCR4 positive cells are shown. **(D-F)** The histogram peaks of CXCR4 staining levels on control or c-MET KO MT-4 cells (D), primary CD4^+^ T cells (E), and MT-4 cells (F) are shown. The results are representative of three independent experiments performed with biological samples (n=3) or primary CD4^+^ T cells from 3 independent healthy donors.

**Fig. S14.**
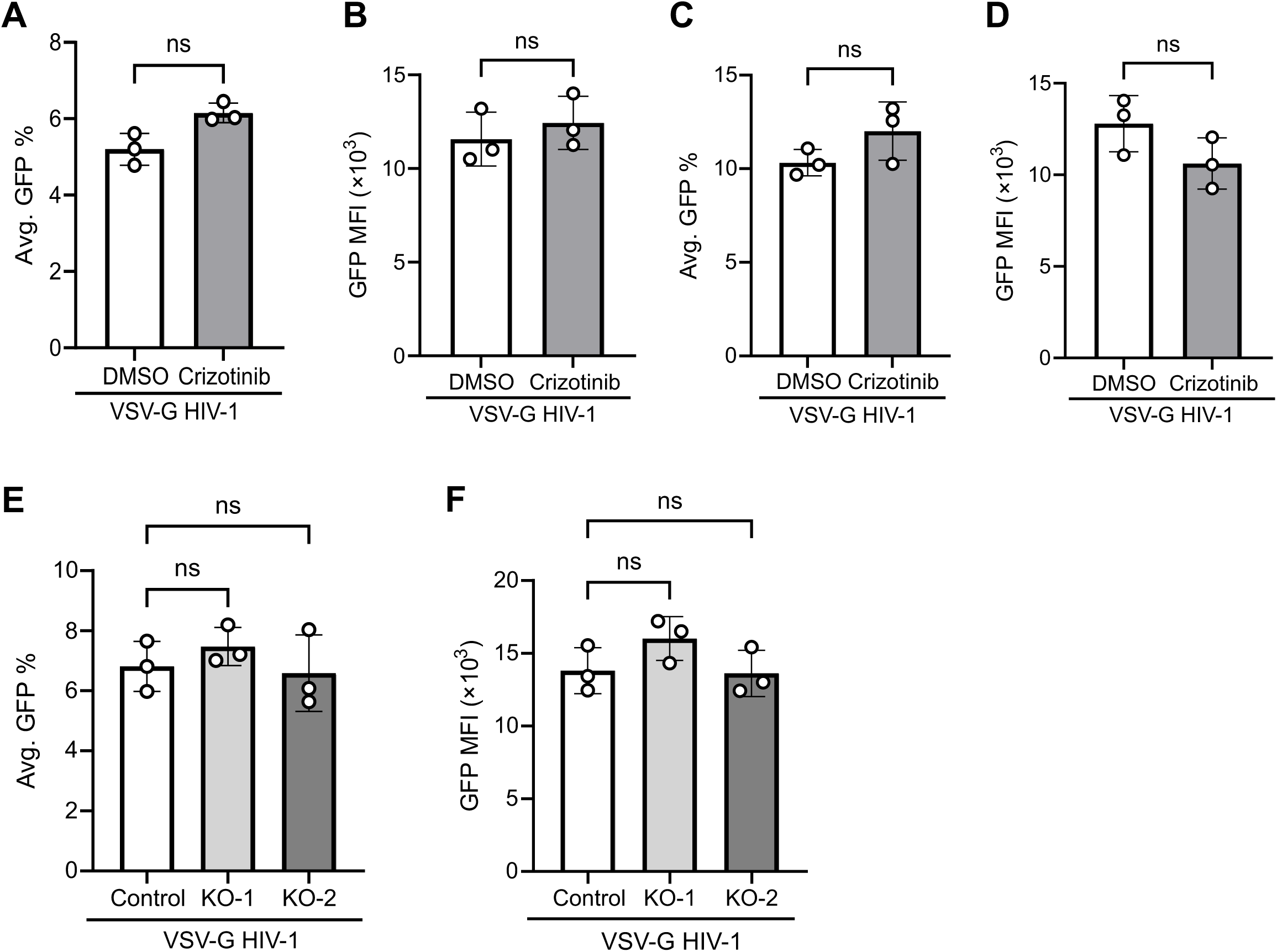
Crizotinib treatment or c-MET KO does not significantly affect VSV-G-pseudotyped HIV-1 infection in CD4^+^ T cells. **(A-F)** VSV-G-pseudotyped HIV-1 infection was measured by GFP expression using flow cytometry at 48 hpi. Changes in average GFP percentage (A) and change in MFI of GFP-positive cells (B) from HIV-1 infected primary CD4^+^ T cells from three independent donors upon DMSO or crizotinib treatment are shown. (C-D) Changes in average GFP percentage (C) and change in MFI of GFP-positive cells (D) from HIV-1 infected MT-4 cells upon DMSO or crizotinib treatment are shown. (E-F) Changes in average GFP percentage (E) and change in MFI of GFP-positive cells (F) from control or c-MET KO MT4 cells followed by HIV-1 infection are shown. ns, not significant.

**Supplementary Excel file information**: Differential expressed genes (DEGs) in control MT-4 cells versus c-MET knockout MT-4 cells (an Excel file). Among these DEGs, 3,458 were significantly upregulated and 2,988 were significantly downregulated in c-MET KO cells as compared with control MT-4 cells.

## Notes

### Competing Interest Statement

The authors have declared no competing interest.

